# Thresholding of sensory inputs by extrasynaptic glutamate receptors in olfactory bulb glomeruli

**DOI:** 10.1101/341958

**Authors:** D.H. Gire, J.D. Zak, J.N. Bourne, N.B. Goodson, B.E. Lynch, C.G. Dulla, N.E. Schoppa

## Abstract

The mammalian olfactory bulb has presented a challenging system for understanding information processing, in part because the bulb largely lacks the topographical ordering of neurons that promotes processes such as lateral inhibition. Here we have used dual and triple-cell recordings in rodent bulb slices combined with ultrastructural methods to provide the first experimental evidence for a processing mechanism circumventing this problem that operates at the level of single glomeruli, the bulb’s odorant receptor-specific modules. A key feature is non-traditional, extrasynaptic glutamatergic signaling derived from excitatory interneurons and what it means for the local balance between excitation (E) and inhibition (I). We found that the distinct dynamic properties of extrasynaptic excitation versus synaptic inhibition create a thresholding effect whereby only strong stimuli produce a favorable E/I balance enabling an output. This single-glomerulus threshold could have a number of important functions during natural odor responses, for example enhancing stimulus tuning.

## Introduction

In many sensory systems, GABAergic interneurons play a central role in sharpening the tuning of excitatory cells, a mechanism that is thought to decorrelate the neural representations for different but similar stimuli (Isaacson and Scanziani, 2011; Franke et al., 2017; Wood et al., 2017). In circuits such as the retina and somatosensory cortex, where neurons are topographically ordered, the GABAergic cells operate mainly through center-surround, lateral inhibitory interactions between nearby, functionally similar cells (Werblin, 1972, 1974; Cook and McReynolds, 1998; Petersen et al., 2003; Adesnik and Scanziani, 2010). The first processing structure in the olfactory system, the olfactory bulb, presents an interesting counter-example. While the bulb appears to be capable of sharpening the tuning of output mitral cells (MCs) and tufted cells (TCs) to odors (Yokoi et al., 1995; Davison and Katz, 2007; Tan et al., 2010), its circuitry is marked by only loose topographical ordering, as reflected in the organization of odorant receptor (OR)-specific glomeruli (Mori et al., 2006; Fantana et al., 2008; Soucy et al., 2009; Ma et al., 2012). This lack of order suggests that the tuning of MC/TCs may be sharpened through “intraglomerular” mechanisms at one glomerulus, either acting alone or in parallel with targeted, long-range lateral inhibition (Economo et al., 2016). Modeling studies have suggested that a single glomerulus could sharpen MC tuning by essentially acting as a threshold that works on the basis of the local balance between excitation and inhibition (the E/I balance; Cleland and Sethupathy, 2006; Linster and Cleland, 2009; Cleland and Linster, 2012): An odor with a low affinity for an OR (a weak stimulus) would drive more inhibition than excitation, thus failing to excite MCs, but a higher affinity odor could drive enough excitation to overcome inhibition. Experimental evidence consistent with such a threshold has come from *in vivo* studies showing a sign-inversion in the MC/TC response from inhibition to excitation with increasing odor concentration (Fukunaga et al., 2014; Economo et al., 2016), a manipulation similar at the level of an OR to increasing odor affinity. There remains however no direct experimental evidence indicating that a single glomerulus can function as a threshold nor what the potential underlying mechanisms could be.

Glomeruli have a number of cell-types that in principle could be part of such a threshold (see **Fig. 1B1**). Besides output MCs and TCs, each glomerulus has two classes of interneurons with apical dendritic arbors confined to one glomerulus, GABAergic periglomerular cells (PG cells) and glutamatergic external tufted cells (eTCs; notably, distinct from the output TCs). eTCs have a particularly interesting position in the circuit in that they are an intermediary in a multi-step, feedforward path of activating MCs by OSNs that exists in parallel with the direct OSN-to-MC pathway (De Saint Jan et al., 2009; Najac et al., 2011; Gire et al., 2012; Vaaga and Westbrook, 2016). The available evidence suggests that the feedforward pathway underlies most of the MC excitatory response to OSN stimulation. eTCs also excite most (∼70% of) PG cells through glutamatergic dendrodendritic synaptic contacts (Hayar et al., 2004a; Shao et al., 2009), and also are a major recipient of inhibition from PG cells (Murphy et al., 2005). Hence, a local network of eTCs and PG cells lies between OSNs and MCs. One intriguing aspect of the glomerular circuit is the mechanism by which eTCs excite MCs. eTCs form few if any direct synaptic contacts onto MCs (Pinching and Powell, 1971; Bourne and Schoppa, 2017), and instead appear to excite MCs through activation of “extrasynaptic” glutamate receptors (Gire et al., 2012). Extrasynaptic transmission, which is likely promoted by confined spaces intrinsic to glomeruli (Rossi et al., 1995; Kinney et al., 1997; Chao et al., 1997; Kasowski et al., 1999), could provide a potentially powerful tool for generating an intraglomerular threshold. In contrast to synaptic glutamate receptors that sense a high glutamate concentration even when a presynaptic cell fires single spikes (Clements et al., 1992), activation of extrasynaptic glutamate receptors generally requires coincident activation of many nearby axons or trains of action potentials to enable accumulation of glutamate at receptor sites (Asztely et al., 1997; Carter and Regehr, 2000; Clark and Cull-Candy, 2002; Chalifoux and Carter, 2011; Coddington et al., 2013; Nahir and Jahr, 2013; Wild et al., 2015). Such dynamics in extrasynaptic glutamate could contribute to an E/I balance for MCs/TCs that favors inhibition over excitation in the presence of a weak stimulus that drives few eTC spikes (e.g., a low-affinity odor) but a much more favorable E/I balance with a strong stimulus.

**Figure 1.**
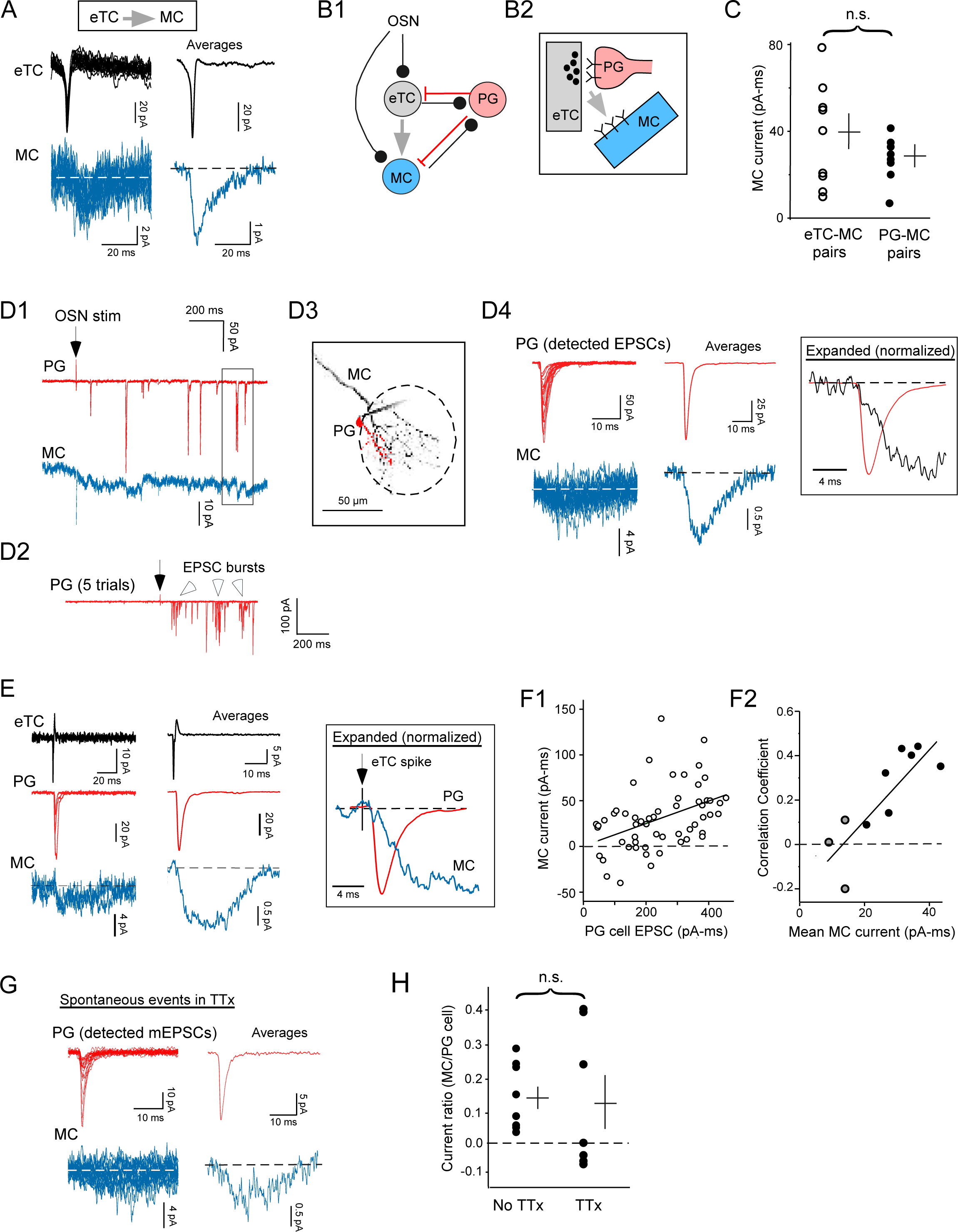
eTC-to-MC transmission involves glutamate spillover at eTC-to-PG cell dendrodendritic synapses. (**A**) Recordings from an eTC-MC pair showing MC currents (dark blue; at *V_hold_* = –77 mV) evoked by single eTC spikes (black; in loose cell-attached, LCA, mode). Raw traces and averages (*n* = 94) are illustrated. (**B1**) Simplified circuitry at a glomerulus: OSN axons contact eTCs (black lines), which in turn send glutamatergic extrasynaptic signals to MCs (gray arrow). eTCs and MCs can excite GABAergic periglomerular cells (PG) at dendrodendritic synapses, which feed back inhibition onto these cells (red lines). The OSN-to-eTC-to-MC feedforward pathway occurs in parallel with the direct OSN-to-MC pathway (left). Not shown: OSN-to-PG cell synapses, which can excite a minority (30%) of PG cells, GABAergic short-axon cells, and other classes of tufted cells besides eTCs. (**B2**) The “spillover” hypothesis: Glutamate released at an eTC-to-PG cell synapse activates extrasynaptic receptors on nearby MC apical dendrites. (**C**) Summary of MC current measurements obtained in 9 eTC-MC pair recordings, as in **A**, and in 8 PG cell-MC pair recordings, as in **D**. Lines reflect mean ± SEM. Charge values, obtained by integrating the inward MC currents, were multiplied by –1. (**D**) Whole-cell current recordings in a same-glomerulus PG cell-MC pair (*V_hold_* = –77 mV in both cells) in response to OSN stimulation (at arrow; 40 A) used to test the spillover hypothesis. Shown are currents associated with a single response-trial (**D1**; PG cell in red), 5 superimposed trials for the PG cell on a less expanded scale (**D2**), a fluorescent image of the pair (**D3**; glomerulus demarcated by dashed oval), and raw examples and average traces (*n* = 67) that reflect detected rapid EPSCs in the PG cell and time-locked MC currents (**D4**). Boxed region in **D1** shows two examples of current deflections in the MC that were time-locked to rapid EPSCs in the PG cell. Open arrowheads in **D2** point to bursts of EPSCs in the PG cell that delineate the cell as the subtype that receives direct input from eTCs (Shao et al., 2009). (**E**) Current events collected from a different, triple-cell recording that included a same-glomerulus eTC, PG cell, and MC. Expanded and normalized average traces (*n* = 78) at far right (boxed) indicate that the PG cell and MC currents both occurred with a 1-2 onset delay after the eTC spike. (**F**) Correlation between individual PG cell and MC current events. Results from the experiment in part **D** (**F1**; correlation coefficient = 0.43, *p* = 0.0010) and a summary across 10 PG cell-MC pair recordings (**F2**) are shown. In the summary, values for the correlation coefficients are plotted as a function of the mean MC current in the same experiment to show that these two parameters were themselves highly correlated (*r* = 0.78, *p* = 0.008). Plot combines data from our standard recordings (black circles) as well as 3 recordings in tetrodotoxin, TTx (see **G**; gray circles). Pair recordings in which no time-locked MC currents were observed were excluded. (**G**) Spontaneous currents in a PG cell-MC pair recorded in TTx (1 μM). Averages reflect 28 events. The presence of MC currents locked to PG cell mEPSCs is direct evidence for spillover at single release sites. (**H**) Ratios of the MC versus PG cell currents (integrated charge for both) in the absence (*n* = 8) and presence of TTx (*n* = 7). The similar ratios under the two conditions argue that spillover accounts for most of eTC-to-MC extrasynaptic transmission (see text).

In this study, we have used a combination of patch-clamp recordings and ultrastructural approaches to investigate extrasynaptic glutamatergic signaling within a glomerulus and its potential role in generating an intraglomerular threshold. Recordings were conducted in rodent olfactory bulb slices rather than *in vivo*, based on the fact that studying single-glomerulus mechanisms in isolation is virtually impossible during odor-evoked responses due to the inherently complex nature of an odor stimulus (Yang et al., 1998; Rubin and Katz, 1999; Uchida et al., 2000). Here we isolated single-glomerulus mechanisms both by performing pair and triple-cell recordings between cells affiliated with the same glomerulus, and also by using strategies to stimulate sensory inputs that restricted neural activity to one glomerulus. As a first step in our study, we better defined the mechanistic underpinnings of extrasynaptic signaling from eTCs to MCs, wherein we provide evidence that it is due to “spillover” of glutamate released at eTC-to-PG cell synapses. This in turn formed the basis for the rest of the study that examined how extrasynaptic glutamatergic signaling in the eTC-MC network contributes to the local E/I balance under various conditions of stimulus strength. Quantitative relationships between stimulus strength and E/I balance were established using two approaches, either in different pair-cell recording combinations, in which we related the number of spikes in an eTC to the excitatory current in excitatory and inhibitory cells, or by using recordings of excitatory and inhibitory currents in response to measurable levels of OSN input. Our results indicated that the distinct dynamics that are intrinsic to extrasynaptic excitation versus synaptic inhibition contribute to a potent intraglomerular thresholding mechanism.

## Results

**Fig. 1A** illustrates a pair-cell recording between an eTC and MC that depicts feedforward MC excitation elicited by single eTC spikes (De Saint Jan et al., 2009; Najac et al., 2011; Gire et al., 2012). That these current signals are extrasynaptic is based on morphological evidence that there are few if any direct synaptic connections between eTCs and MCs (Pinching and Powell, 1971; Bourne and Schoppa, 2017), combined with the relatively small size and slow kinetics of the MC currents (integrated charge = –39 ± 8 pA-ms, 20-80% rise-time = 3.6 ± 0.5 ms, half-width = 27 ± 4 ms, *n* = 9; **Fig. 1C**). In addition, eTC-to-MC currents are selectively suppressed by a low-affinity AMPA receptor antagonist γ-d-glutamyl glycine (Gire et al., 2012). Such an antagonist should act preferentially at extrasynaptic receptors at which the glutamate concentration is relatively low. In this study, we first investigated the mechanisms underlying eTC-to-MC extrasynaptic signaling, which then formed the basis for the rest of the study that investigated its contribution to the local excitation/inhibition (E/I) balance within a glomerulus.

### Extrasynaptic eTC-to-MC excitation is due to spillover at eTC-to-PG cell synapses

Our preliminary hypothesis for what underlies extrasynaptic eTC-to-MC excitation involved the well-established dendrodendritic excitatory synapses from eTCs onto GABAergic PG cells (Hayar et al., 2004a; **Fig. 1B1,2**). Glutamate released at these synapses could directly activate postsynaptic glutamate receptors on PG cells and also diffuse (i.e., spill over) out of the synaptic cleft and activate nearby glutamate receptors on MC apical dendrites (Salin et al., 2001). To test the spillover hypothesis, simultaneous voltage-clamp recordings were performed in cell-pairs that included a MC and a PG cell affiliated with the same glomerulus (*V_hold_* = –77 mV in both cells; **Fig. 1D1,3**). The presence of excitatory currents in MCs that were precisely time-locked to rapid AMPA receptor-mediated excitatory post-synaptic currents (EPSCs) in PG cells would provide evidence for spillover. All of the PG cells analyzed displayed complex current responses that included bursts of EPSCs (**Fig. 1D2**) that are characteristic of the ∼70% of PG cells excited by eTCs (Hayar et al., 2004a; Shao et al., 2009), but we focused here on isolated EPSC events in the PG cells (≥ 50 ms event separation) because of the requirement for precise temporal information in our analysis (see *Analysis* section of *Methods* for additional details).

We found clear evidence for potential spillover, both during responses to electrical stimulation of olfactory sensory neurons (OSNs; 100 μsec pulse at 5-44 μA; 5 pairs, including example in **Fig. 1D**) as well as during recordings of spontaneous currents (3 pairs). Seven of the 8 PG cell-MC pairs displayed small currents in the MC that were locked to the rapid EPSCs in PG cells (mean onset latency between averaged PG cell and MC currents = 0.2 ± 0.1 ms, *n* = 7; **Fig. 1D4**). The MC currents in the PG cell-MC pairs moreover closely resembled MC currents evoked by single eTC spikes in eTC-MC pairs (**Fig. 1A**), both in their magnitude (integrated charge = –28 ± 5 pA-ms, *n* = 8; **Fig. 1C**) and in their slow kinetics (20-80% rise-time = 3.3 ± 0.5 ms, half-width = 29 ± 7 ms, *n* = 7; *p* > 0.10 in Mann-Whitney U test comparing all variables). In the PG cell-MC pair recordings, we could not be certain that all of the time-locked currents reflected glutamate release from eTCs, but direct evidence that eTCs drove coupled currents was obtained in two (out of 10) triple-cell recordings that included MCs, PG cells, and eTCs at the same glomerulus (example in **Fig. 1E**). In these, the MCs displayed small currents (mean integrated charge = –42 and –20 pA-ms) that were time-locked to both rapid EPSCs in the PG cell and spikes in the eTC.

While the MC currents time-locked to PG cell EPSCs were consistent with spillover at eTC-to-PG synapses, another possibility is that they reflected glutamate release at “ectopic” release sites, i.e., sites with no dedicated post-synaptic partners (Matsui and Jahr, 2003; Coggan et al., 2005). A spike in an eTC could have resulted in glutamate release at eTC-to-PG cell synapses and nearly coincident release at a different, ectopic site. However, arguing that release occurred at the same sites was the fact that, within each pair-cell recording, the MC current events were correlated in size to their associated PG cell EPSCs (mean correlation coefficient *r* = 0.30 ± 0.03, *n* = 7; *p* ≤ 0.014 in 5 of 7 pairs with coupled currents; **Fig. 1F1**). Such correlations would arise if the MCs and PG cells were responding to the same bolus(es) of glutamate, which should naturally vary in magnitude due to variability in the amount of glutamate in a vesicle and/or the number of vesicles released per action potential. The correlation coefficients were modest, but this could be explained by the small size and low signal-to-noise of the MC currents. Indeed, consistent with this explanation, we found that the values of the correlation coefficient across the pairs were themselves positively correlated to the mean MC current in the pairs (*r* = 0.78, *p* = 0.008; **Fig. 1F2**). Additional, more direct evidence for spillover was obtained in PG cell-MC pair recordings conducted in the presence of the sodium channel blocker tetrodotoxin (TTx, 1 μM; **Fig. 1G).** In 3 of 7 such recordings, small MC currents (mean integrated charge = –12 ± 2 pA-ms, *n* = 3) were observed that were time-locked to spontaneous miniature EPSCs (mEPSCs) in PG cells (mean peak amplitude = –12 ± 1 pA, mean current onset-latency between PG cell and MC currents = 0.4 ± 0.3 ms, *n* = 3). Because spontaneous glutamate release in TTx should generally be occurring at single release sites at a time, the slow MC currents locked to the mEPSCs had to result from glutamate release at the same sites.

Could ectopic glutamate release contribute to a large component of eTC-to-MC transmission even with the clear evidence for discrete, single-site spillover events in TTx? The PG cell-MC pair recordings conducted in TTx provided one other useful measure that enabled us to address this question. This involved comparing the *ratio* of the MC and PG cell currents (the MC/PG cell current ratio) recorded in the presence versus absence of TTx. Because ectopic release sites should contribute to spike-evoked MC currents locked to EPSCs in PG cells but not single-site MC currents locked to mEPSCs, the MC/PG cell current ratio should have been much less in TTx if ectopic glutamate release was more dominant than spillover. However, we found that the current ratios under the two conditions were indistinguishable (**Fig. 1H**; No TTx: 0.14 ± 0.04, *n* = 8; TTx: 0.13 ± 0.09, *n* = 7*; p* > 0.10 in Mann-Whitney U test; analysis included all recordings, including those in which no time-locked MC current was evident). Thus, spillover and not ectopic release appeared to contribute to most of eTC-to-MC extrasynaptic transmission.

As a final, independent check on the spillover hypothesis, ultrastructural studies were conducted on olfactory bulb slices in which eTCs were labeled with biocytin during whole-cell patch recordings (Bourne and Schoppa, 2017; **Fig. 2**). In reconstructions based on electron micrographs, we found ample evidence for complexes that could underlie spillover at eTC-to-PG cell synapses. Amongst 13 identified dendrodendritic synapses from eTC cells onto inhibitory dendrites (mostly presumed to be PG cells), 11 had nearby glutamatergic dendrites (within 0.5 μm) that were presumed to be MCs (see *Methods;* two examples in **Fig. 2A1-3** and **2B1-3; Fig. 2D**). Furthermore, glial processes could be identified that extended close to the eTC-PG cell-MC complexes (in 5 of 13 synapses) and sometimes surrounded the complexes (**Fig. 2A3**). Importantly, the glial processes never surrounded the eTC-to-PG cell synapse, isolating it from the presumed MC dendrite (*n* = 11). Because these processes and their associated transporters limit glutamate spillover at excitatory synapses (Asztely et al., 1997; Murphy-Royal et al., 2017), their absence would greatly facilitate the ability of glutamate released from eTCs to act on extrasynaptic glutamate receptors on MCs.

**Figure 2.**
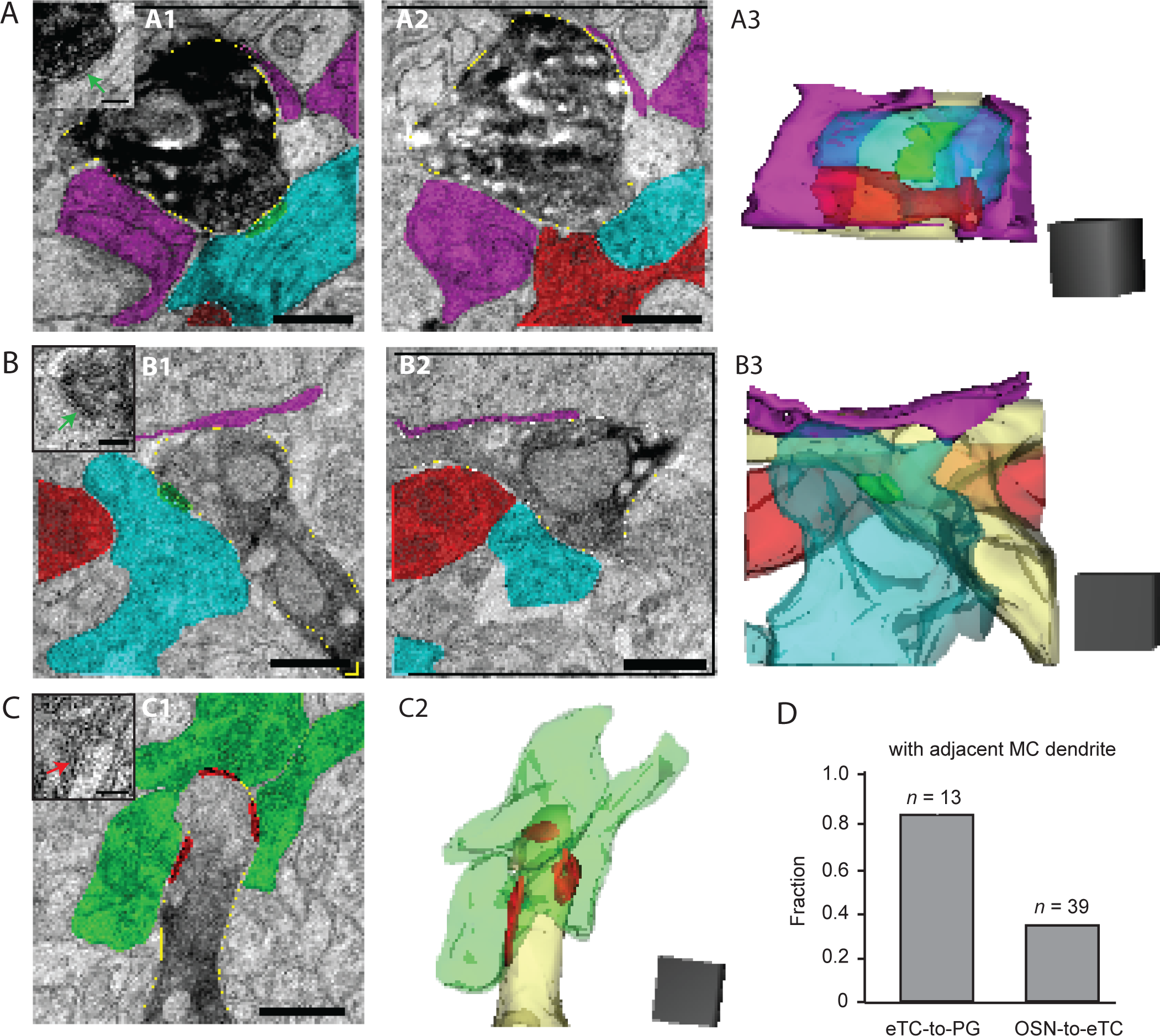
Ultrastructural evidence for complexes that could support spill-over. (**A**) Sequential electron micrographs (**A1** and **A2**) and three-dimensional reconstruction (**A3**) of an example complex that includes a DAB-labeled eTC dendrite (darkened in micrographs; light yellow in reconstruction) forming a synapse (green) onto a putative PG cell dendrite (blue). A putative MC dendrite (red) and glial processes (purple) are in close proximity. Note that the glial processes in the reconstruction appear to surround the dendrites. Inset in part **A1** shows the same image as in **A1** but without the colors so that the synapse (green arrow) can be seen more clearly. Scale bars: 0.5 microns for micrographs, 0.1 microns for inset of **A1**. Scale cube in **A3** = 0.5 μm^3^. (**B**) Another example of a complex containing an eTC-to-PG cell synapse with adjacent glial process and putative MC dendrite. Note that in the reconstruction the putative MC dendrite runs behind the PG cell and eTC dendrites. Scale bars and cube as in part **A**. (**C**) Electron micrograph (**C1**) and reconstruction (**C2**) of a cluster of synapses (in red) from OSNs (green) onto a DAB-labeled eTC dendrite (light yellow in reconstruction). Note the absence of an adjacent MC dendrite. Inset in **C1**: Original image of the eTC dendrite showing one of the OSN synapses (red arrow). Scale bars and cube as in part **A**. (**D**) Summary of analysis of 13 eTC-to-PG cell synapses and 39 OSN-to-eTC synapses (pooled results from analysis of two eTC fills in two bulb slices). The fraction out of the total with an adjacent MC dendrite (within 0.5 μm) was much higher for eTC-to-PG cell synapses.

### Absence of spillover at OSN synapses

Because glomeruli in the bulb are compartmentalized structures with confined spaces (Chao et al., 1997; Kasowski et al., 1999), we wondered whether glutamate spillover is restricted to eTC-to-PG cell dendrodendritic excitatory synapses. Spillover of glutamate at OSN synapses onto eTCs (**Fig. 3A**) in particular could, like spillover at eTC-to-PG synapses, impact information flow from OSNs to MCs. We thus also performed simultaneous voltage-clamp recordings in eTC-MC pairs (*V_hold_* = –77 mV in both cells; **Fig. 3B1,B2**), examining whether MCs displayed slow currents time-locked to fast EPSCs in eTCs. Fast EPSCs that reflected OSN-to-eTC transmission (“OSN-EPSCs”) could be easily identified as spontaneous events in the eTC recordings (Hayar et al., 2005). Interestingly, MCs in the eTC-MC pairs did not display currents coupled to OSN-EPSCs in eTCs (–0.2 ± 0.6 pA-ms integrated charge, *n* = 6; **Fig. 3B3,C,D)**, inconsistent with spillover. This conclusion was also supported by our ultrastructural data (**Fig. 2C1,C2,D**). In contrast to the eTC-to-PG cell synapses (**Fig. 2A,B,D**), which had presumed MC dendrites in close proximity in the large majority (85%) of examples examined, OSN synapses onto labeled eTCs had nearby dendrites of presumed MCs relatively infrequently (38%; 15 out of 39 OSN-to-eTC synapses).

**Figure 3.**
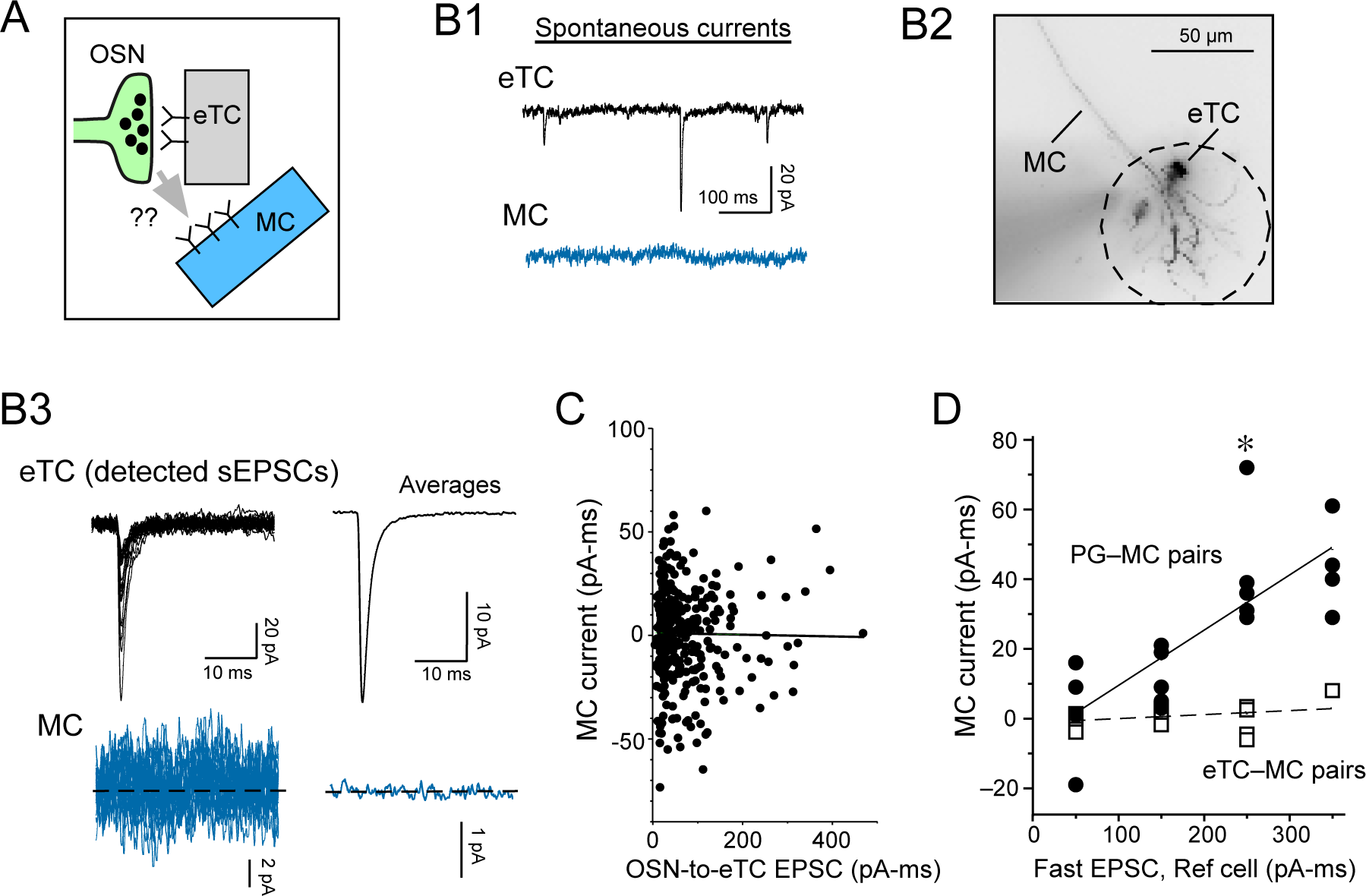
Absence of spillover at OSN synapses. **(A)** Model to be tested: Glutamate released at OSN-to-eTC synapses spills over and activates glutamate receptors on adjacent MC dendrites. **(B)** Example whole-cell recording from an eTC-MC pair at the same glomerulus used to test for spillover at OSN-to-eTC synapses. Raw traces recorded without a stimulus **(B1),** an image of the pair (**B2;** collapsed in the z-axis), and examples and averages (*n =* 293) of detected spontaneous OSN-EPSCs (sEPSCs) in the eTC and time-locked MC currents **(B3)** are illustrated. The cell body of the eTC in this pair was just above the glomerulus to which the MC sent its apical dendrite. **(C)** X-Y plot relating the magnitude of the OSN-EPSCs and associated MC currents for the experiment in **B.** Line reflects linear regression fit of the data (*r* = −0.012; *p* = 0.82). **(D)** Summary of MC currents recorded in eTC-MC pairs (open squares) versus PG cell-MC pairs (filled circles), plotted as a function of the amplitude of the fast EPSC in the reference cell (either eTC or PG cell). Data were binned according to the magnitude of the fast EPSCs in 100 pA-ms increments. Note that, while the MC current in the PG cell-MC pairs clearly increased as a function of the magnitude of the PG cell EPSC (linear regression fit: *r =* 0.79, *p* = 0.000056), no such relationship was observed in the eTC-MC pairs (*r* = 0.28, *p* = 0.31). The eTC-MC pair dataset includes 4-5 points each in the bins centered at 50, 150, and 250 pA. Each point in each bin reflects a single experiment. \**p* = 0.01 in Mann-Whitney U test comparing the MC currents in the two cell-pair types for similar-amplitude fast EPSCs in the reference cell.

Amongst the potential caveats to our physiological results arguing against spillover at OSN synapses was the fact that the OSN-EPSCs in eTCs in the eTC-MC pairs were relatively small, with typical magnitudes near 20 pA (mean peak amplitude = –21 ± 3 pA, *n* = 6). This compared to values that were ∼3-fold larger for the eTC-to-PG cell EPSCs in the PG cell-MC pairs (mean peak amplitude = –63 ± 15 pA, *n* = 8; *p* = 0.028 in unpaired t-test). The small OSN-EPSCs in eTCs suggested that the absence of coupled MC currents was because OSN-to-eTC synapses were relatively weak, making it difficult to detect spillover. However, we found no coupled MC currents in the eTC-MC pairs (–1 ± 2 pA-ms integrated charge, *n* = 4) even when we focused on a subset of larger OSN-EPSCs (200-300 pA-ms; *p* = 0.01 in Mann-Whitney U test comparing MC currents coupled to eTC versus PG cell EPSCs of the same amplitude; **Fig. 3D**). The absence of coincident current events in the eTC-MC pairs was also not because the cells were affiliated with different glomeruli. In these experiments, we used two independent criteria to confirm that the eTC-MC pairs were affiliated with the same glomerulus, including the cells’ anatomy (**Fig. 3B2**) as well as the co-occurrence in the current recordings of discrete, large-amplitude events associated with long-lasting depolarizations (LLDs; **Supplementary Fig. 1**). LLDs are well-known to co-occur in all MCs and eTCs at the same glomerulus (Carlson et al., 2000; Gire and Schoppa, 2009). Thus, we take the absence of coupled currents in the eTC-MC pair recordings as strong evidence against spillover at OSN-to-eTC synapses.

In the present study, we did not explicity test for spillover at other glomerular synapses not involved in feedforward signaling from OSNs to MCs, for example the possibility of MC-to-MC signaling via spillover of glutamate released at MC synapses onto PG cells. Prior studies have provided evidence for extrasynaptic signaling between MCs (Isaacson, 1999; Schoppa and Westbrook, 2002; Pimentel and Margrie, 2008), although spillover has not been tested.

### Excitation of excitatory and inhibitory cells as a function of increasing eTC spike number

Having established that a major part of extrasynaptic glutamatergic transmission from eTCs onto MCs is mediated by spillover at eTC-to-PG cell synapses, we next turned to the functional question of what such a mechanism means for the balance between excitation and inhibition (the E/I balance) within a glomerulus. Modeling studies have suggested that changes in the local E/I balance as a function of the strength of a stimulus could underlie a thresholding mechanism that enables selective activation of MC/TCs when OSN inputs are strong (e.g., in the presence of odors with a high OR affinity; Cleland and Sethupathy, 2006; Linster and Cleland, 2009; Cleland and Linster, 2012). As a first step, a series of pair-cell recordings were performed in which excitatory currents were measured in either MCs or PG cells as a function of the number of spikes in an eTC. This reduced system enabled us to establish quantitative relationships between one measure of stimulus strength (eTC spike number) and the magnitude of excitation of the excitatory or inhibitory cells in the local glomerular network. For the analysis of extrasynaptic excitation, we augmented our data set comprised of eTC-MC pairs with additional recordings in cell pairs in which both participants were eTCs. The properties of extrasynaptic excitation in eTC-eTC pairs (see example recording in **Fig. 4F**) could be more directly linked to the analysis of eTC responses to OSN stimulation conducted later in the study (see **Figs. 8-9**). In the various pair-cell recordings, the eTCs were induced to spike with direct current injection (during whole-cell recordings; example in **Fig. 4A**) or we took advantage of the natural tendency of eTCs to engage in spontaneous spike bursts of varying number (Hayar et al., 2004b) while eTCs were patched in loose cell-attached (LCA) mode (examples in **Fig. 4B** and **Fig. 5A**).

**Figure 4.**
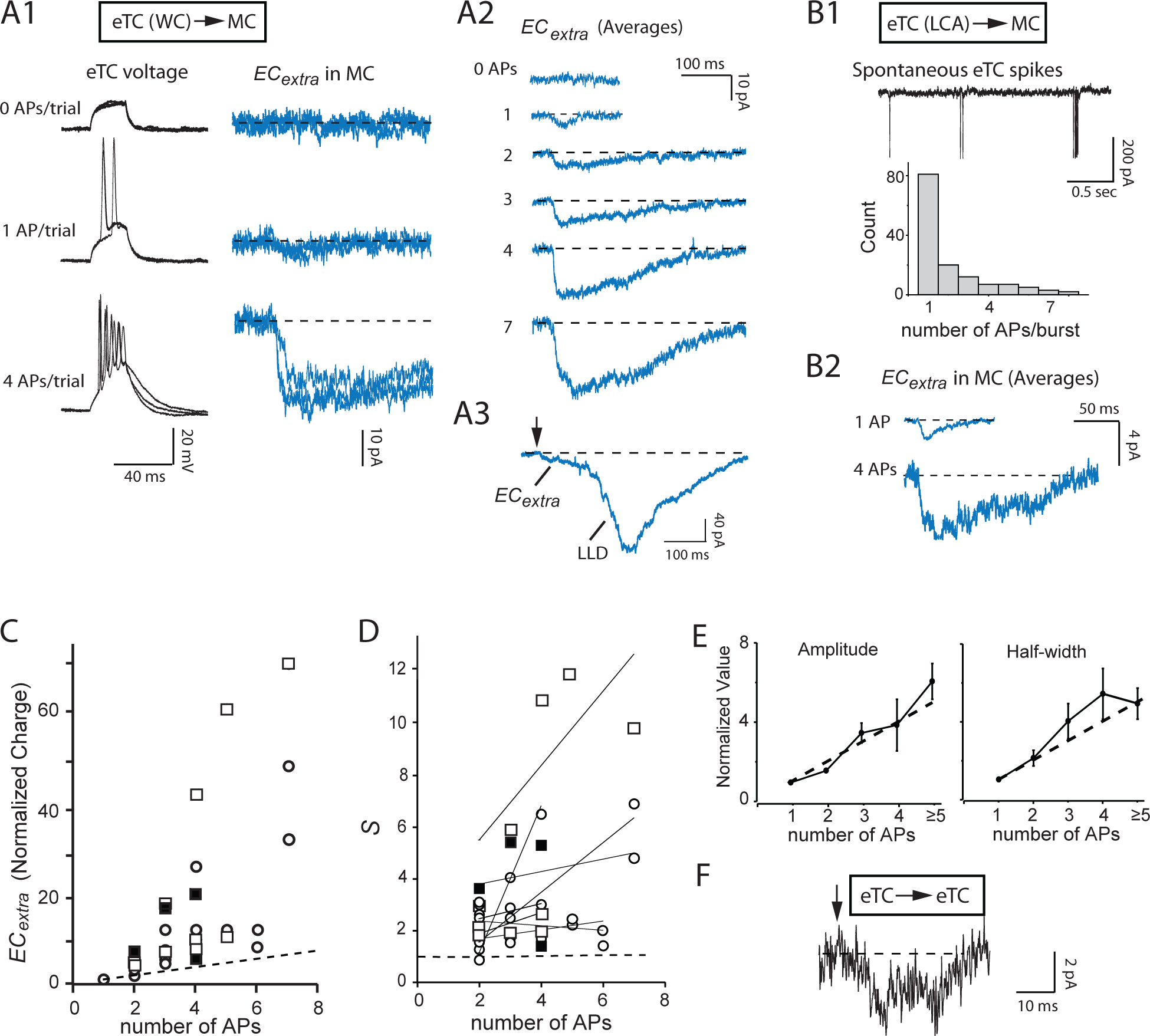
Supralinear increase in extrasynaptic excitatory currents (*EC_extra_*) (**A**) Example whole-cell recordings from an eTC-MC pair illustrating MC current responses to different numbers of action potentials (APs) in the eTC. eTC spikes were evoked by direct depolarizing current pulses (500 pA, 25 ms). Note in the raw traces (**A1**; 3 superimposed trials) and averages (**A2**) that the total current associated with *EC_extra_* for 2 eTC spikes was much larger than linear summation of the 1-AP current. In one of the trials in which the eTC spiked 7 times in the same experiment (**A3**), the MC response included a distinct, much larger LLD event ∼150 ms after the onset of *EC_extra_* (note scale bar). (**B**) Another example eTC-MC pair recording in which the number of spikes in the eTC recorded in the LCA mode (**B1**) was related to *EC_extra_* in the MC (**B2**). As is clear from the raw trace and histogram in **B1**, the eTC spontaneously engaged in both isolated single spikes and bursts of spikes of varying number (Hayar et al., 2004a). (**C**) Summary of *EC_extra_* measurements from 9 eTC-MC pair recordings (open symbols) versus number of spikes in the eTC. Values, which reflect experiments in which the eTCs were in whole-cell (open squares) or LCA (open circles) patch modes, were normalized to the charge elicited by one eTC spike in the same pair recording. Note that nearly all points lie above the dashed line, which reflects linear summation of *EC_extra_* elicited by one eTC spike. Black squares reflect a similar analysis of *EC_extra_* in 4 eTC-eTC pair recordings (see part **F**). Most eTC-MC pair recordings contributed multiple data points to the plot. (**D**) The same data as in part **C**, but with normalized charge measurements converted to estimates (*S*) of the deviation from linearity (see *Methods*; linear: *S* = 1). Linear regression fits of data from 8 pair-cell recordings in which at least three eTC spike numbers ≥ 2 were sampled, indicated that *S* values were themselves generally positively correlated to spike number. (**E**) The supralinear increase in *EC_extra_* was due to increases in both the amplitude (left) and duration (right) of the current. Dashed lines reflect linearity for each parameter. (**F**) Example excitatory current in an eTC evoked by single spikes in another eTC (at arrow) in an eTC-eTC pair recording (average of 12 trials).

**Figure 5.**
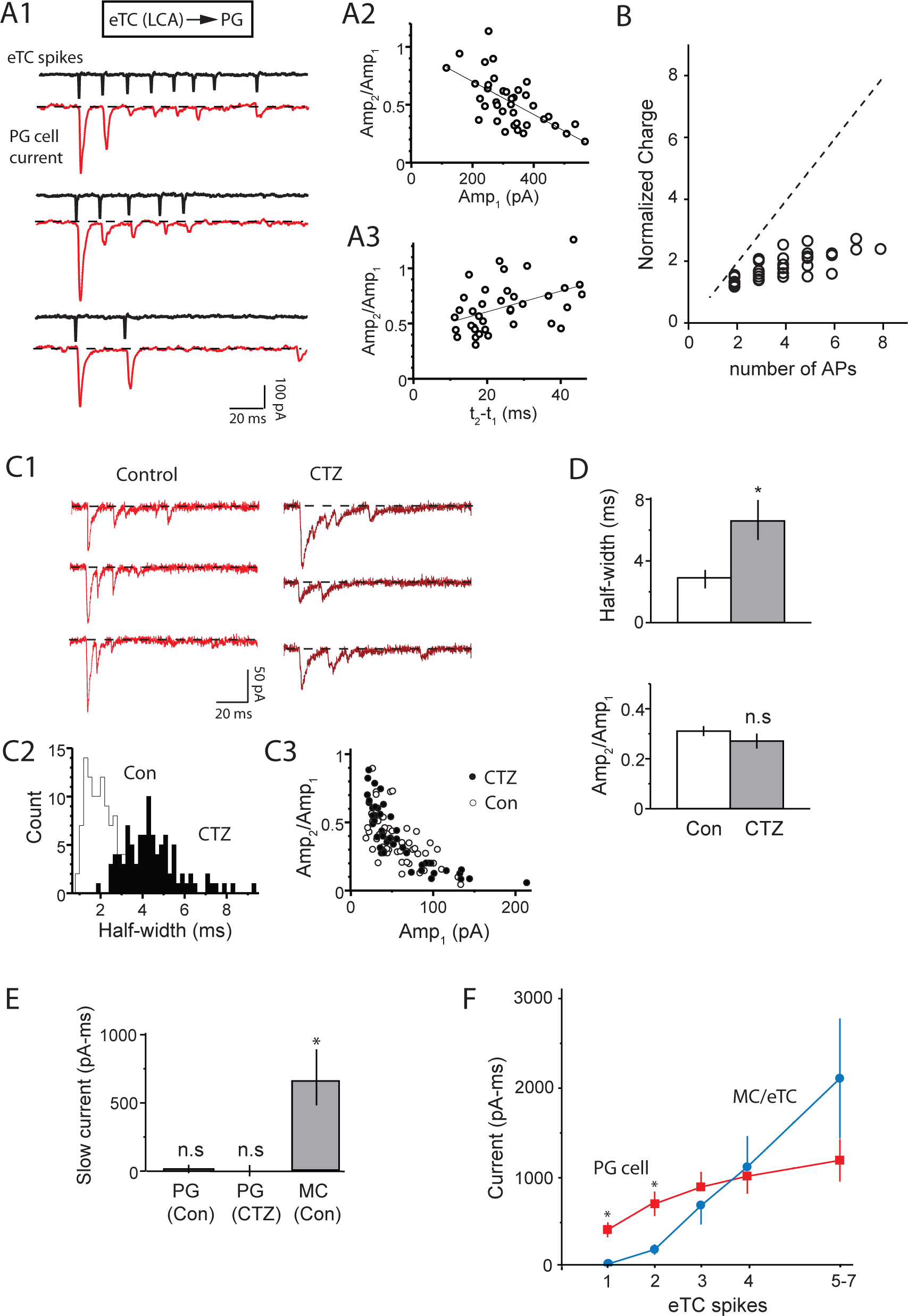
Sublinear increase in PG cell excitatory currents in eTC-PG cell pairs. (**A**) Example recordings in an eTC-PG cell pair showing that spontaneous bursts of spikes in the eTC (recorded in LCA mode) were associated with strongly depressing rapid EPSCs in the PG cell. Three selected examples with varying number of eTC spikes are illustrated in **A1**. Analysis of 37 such bursts in this pair indicated that the degree of synaptic depression as reflected in the amplitude ratio of the first two EPSCs in the burst (*Amp_2_/Amp_1_*) was negatively correlated with the first EPSC amplitude (*r* = –0.63, *p* < 0.0001; **A2**) and positively correlated with the interval between the first two EPSCs (*r* = 0.44, *p* = 0.0058; **A3**). Both are consistent with a presynaptic vesicle depletion mechanism for depression. (**B**) Summary of integrated current measurements from 10 eTC-PG cell pair recordings, demonstrating that the synaptic depression resulted in a sublinear increase in the excitatory current as a function of spike number in the eTC. Diagonal line reflects linearity. (**C**) Example PG cell recording showing that the AMPA receptor allosteric modulator cyclothiazide (CTZ, 100 M) prolonged the EPSCs without altering the degree of depression. Selected EPSC bursts under each condition are illustrated in **C1**, while EPSC half-widths (**C2**) and amplitude ratios (*Amp_2_/Amp_1_*; **C3**) for the same experiment are also shown. The absence of an effect of CTZ on depression supported a presynaptic mechanism. (**D**) Histograms summarizing CTZ effects on EPSC half-width (top) and the *Amp_2_/Amp_1_* ratio (bottom) from 5 PG cell recordings. \**p* = 0.020 in paired t-test. (**E**) Estimates of the magnitude of slow currents in PG cells not directly associated with rapid EPSCs. PG cell currents, both under control conditions and in CTZ, were measured in a 100-200 ms window after the start of each eTC spike burst for bursts with 4 spikes lasting less than 100 ms. For comparison, MC currents (reflecting *EC_extra_*) measured in the same manner are also plotted. *n* = 5 for each recording type. \**p* < 0.01 in paired t-test comparison with zero current. (**F**) Summary of absolute charge measurements as a function of eTC spike number from all pair-cell recordings: eTC-MC or eTC-eTC pairs (blue), eTC-PG cell pairs (red). Each data point reflects mean ± SEM from 7-10 recordings. \**p* < 0.01 in Mann-Whitney U test, Bonferroni correction for multiple comparisons.

Across 9 eTC-MC pairs, we found that an increasing number of spikes in the eTC (bursts defined by clusters of spikes each separated by <50 ms) led to a striking increase in the extrasynaptic excitatory current (called *EC_extra_*, in the rest of the study) recorded in the MC (**Fig. 4A1,A2,B2**). The increase in the *EC_extra_*, charge, which reflected increases in both current amplitude and duration (**Fig. 4E**), was far more than would be expected based on the linear summation of currents elicited by single eTC spikes (**Fig. 4C**). Defining a parameter *S* as a measure of the degree of “supralinearity” (see *Methods*; *S* = 1 is linear), we found that *EC_extra_*, in MCs driven by eTC spike bursts (≥ 2 spikes) was on average ∼3-fold larger than that expected from linear summation (*S* = 2.7 ± 0.3, *n* = 30 spike numbers across 9 pairs; *p* < 0.005 in Wilcoxon matched-pairs signed-rank test). Similar results were obtained from the analysis of *EC_extra_*, in eTC-eTC pairs (*S* = 3.9 ± 0.9, *n* = 4 spike numbers across 4 pairs; **Fig. 4C,F**). In addition, across different eTC spike numbers, the degree of deviation from linearity itself increased with increasing spike number (slope of lines fitted to *S* values = 0.75 ± 0.35, *n* = 8; *p* < 0.05 in Wilcoxon matched-pairs signed-rank test; **Fig. 4D**). Importantly, the supralinearities did not simply reflect changes in the probability of the glomerular LLD. Evoked LLDs, which were occasionally observed in these pair-cell recordings when the spiking eTC engaged in at least 4 spikes (**Fig. 4A3**), were easily distinguishable by size and ignored in this quantitative analysis of the MC/eTC currents (see *Discussion*).

While glutamatergic extrasynaptic excitation of MCs and eTCs rose in a highly supralinear fashion with eTC spike number, synaptic excitation of GABAergic PG cells displayed markedly different properties (**Fig. 5**). The bursts of rapid EPSCs in the PG cell that were driven by eTC spike bursts were characterized by strong depression (**Fig. 5A1**; mean ratio of first and second EPSC amplitudes, *Amp_1_/Amp_2_* = 0.32 ± 0.02, *n* = 7), resulting in a sublinear increase in excitation with eTC spike number (*S* = 0.53 ± 0.03, 33 spike numbers from 7 eTC-PG cell pairs; *p* < 0.005 in Wilcoxon matched-pairs signed-rank test; **Fig. 5B).** Such strong depression occurring on a rapid time-scale is most commonly caused by changes in the availability of presynaptic vesicles (Fioravante and Regehr, 2011). Indeed, as expected for such a mechanism, the degree of depression was positively correlated with the first EPSC amplitude (**Fig. 5A2**) and negatively correlated with the time difference between the first and second eTC spike (**Fig. 5A3**). Furthermore, we found that the AMPA receptor allosteric modulator cyclothiazide (CTZ; 50-100 μM), which slowed post-synaptic AMPA receptor desensitization (116 ± 30% increase in EPSC half-width, from 3.0 ± 0.6 ms to 6.8 ± 1.4 ms, *n* = 5; *p* = 0.020 in paired t-test; Trussell et al., 1993), did not alter EPSC depression (mean decrease in *Amp_1_/Amp_2_* = 0.04 ± 0.04, *n* = 5, *p* = 0.30 in paired t-test; **Fig. 5C,D**). If the EPSC depression were due to desensitization of post-synaptic AMPA receptors, CTZ should have reduced the depression. Notably, the PG cells in these recordings displayed no evidence for prolonged extrasynaptic transmission (**Fig. 5E**) that was characteristic of MCs in the eTC-MC pairs (**Fig. 4**). Selected EPSC bursts in PG cells with ≥4 fast EPSCs had no late current component that could be distinguished from the fast EPSCs (mean integrated current 100-200 ms after first EPSC = –10 ± 18 pA-ms, *n* = 5).

To understand how the observed differences in the dynamics of eTC-driven excitation of MCs/eTCs versus PG cells translated to the balance between the two, we computed the absolute (rather than normalized) level of excitatory current in MCs/eTCs versus PG cells as a function of eTC spike number (**Fig. 5F**). When an eTC engaged in only one spike, the synaptic excitatory current in PG cells (–443 ± 83 pA-ms; *n* = 7) was ∼10-fold greater than *EC_extra_* in MCs/eTCs (integrated charge = –42 ± 6 pA-ms, *n* = 13 pairs; *p* < 0.01 in Mann-Whitney U test), but this difference dissipated as eTC spike number increased. This reflected the supra-versus sublinear increases in excitation of MCs/eTCs versus PG cells. These measures of excitatory current in different cell-types suggest that inhibition mediated by PG cells should be favored when eTCs are weakly excited but extrasynaptic excitation should become much more prominent as eTC excitation increases, a prediction that will be further tested below (*see* **Figs. 8** and **9**).

### Mechanisms underlying supralinear increase in EC_extra_

What drove the supralinear increase in *EC_extra_*? One reasonable hypothesis is that it reflected the long distance between the site of glutamate release on the eTC and the extrasynaptic glutamate receptors on MC/eTC dendrites (**Fig. 1B2**). With such an arrangement, glutamate transients sufficient to activate glutamate receptors may require the accumulation of glutamate generated across multiple spikes (Asztely et al., 1997; Carter and Regehr, 2000; Clark and Cull-Candy, 2002; Chalifoux and Carter, 2011; Coddington et al., 2013; Nahir and Jahr, 2013; Wild et al., 2015). We sought to test one prediction of the distant-receptor model, which is that pharmacological blockade of glutamate transporters aligning the extrasynaptic space should enhance extrasynaptic currents induced by spike bursts (Asztely et al., 1997). This is indeed what we found. Across 5 eTC-MC pair recordings, application of the astroglial glutamate uptake inhibitor DL-threo-beta-benzyloxyaspartate (DL-TBOA, 50-100 μM) caused a near doubling of *EC_extra_* induced by eTC spike bursts (119 ± 15% increase in integrated current for bursts of at least 2 spikes; *p* = 0.0013 in paired t-test, Bonferroni correction for multiple comparisons; **Fig. 6A,C**). At the same time, DL-TBOA did not impact synaptic EPSCs in PG cells (0 ± 7% change in integrated current, *n* = 5, *p* = 0.88 in paired t-test**; Fig. 6B,C**) nor *EC_extra_* when the eTC engaged in only single spikes (3 ± 11% increase, *n* = 5, *p* = 0.80 in paired t-test; **Fig. 6A,C**). We interpret the absence of an effect of DL-TBOA on *EC_extra_* evoked by single eTC spikes to reflect the fact that the glutamate transporters are located at a significant distance away from the glutamate release sites such that they impact only larger, more wide-spread glutamate transients. Such a scenario is consistent with our ultrastructural data (**Fig. 2A,B**) that suggest that glial processes are not directly adjacent to eTC-to-PG cell synapses and instead surround eTC-PG cell-MC spillover complexes.

**Figure 6.**
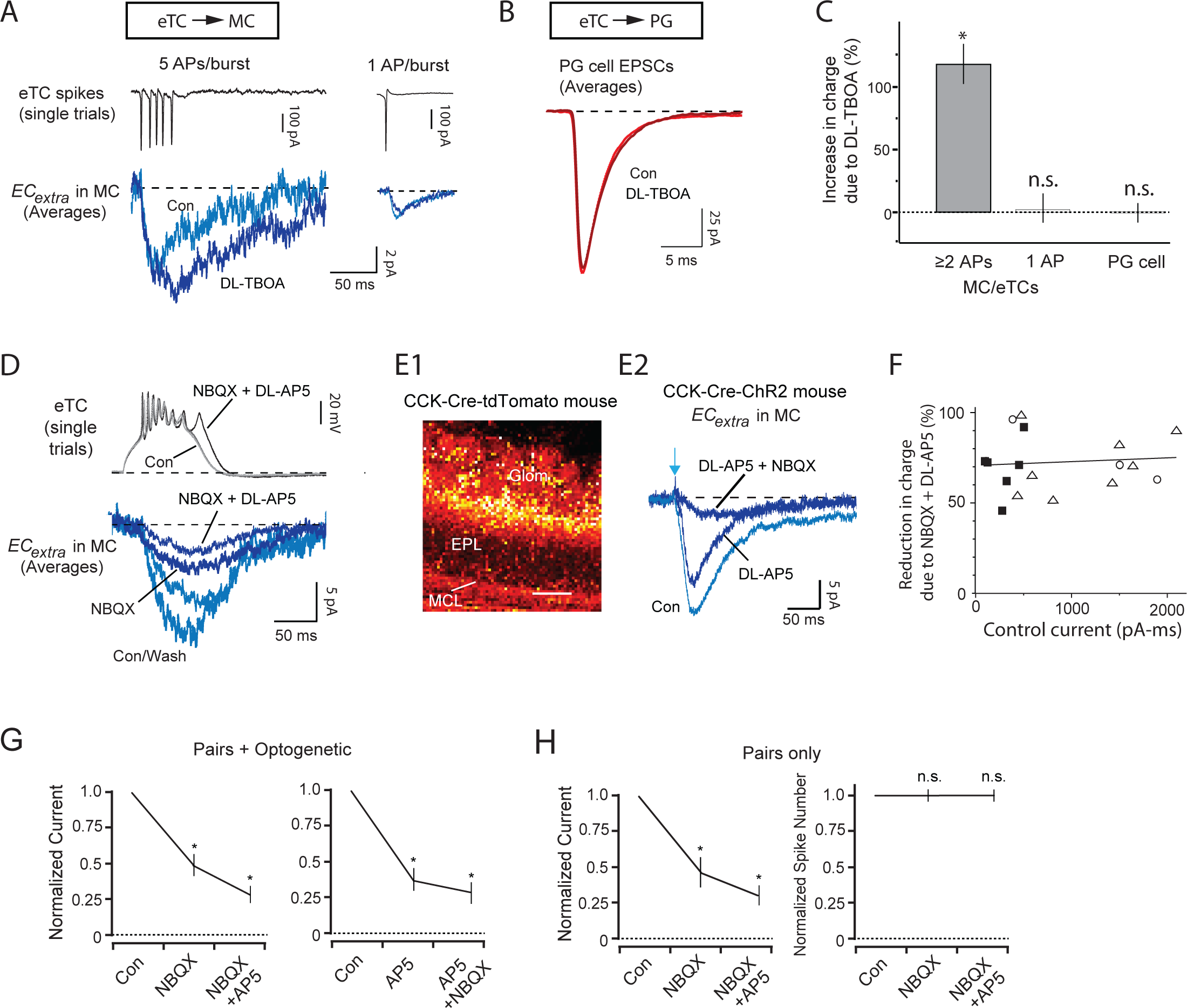
Glutamate receptor-mediated mechanisms underlying supralinear increase in *EC_extra_*. (**A**) Example recordings from eTC-MC pair showing effects of the glial glutamate transporter blocker, DL-TBOA (50 M), on *EC_extra_* in MCs. Note that DL-TBOA enhanced *EC_extra_* when the eTC fired 5 APs (*left*; averages of 12-15 bursts), but not when the eTC spiked only once (*right*; averages of 60). (**B**) Example recording of rapid EPSCs in a PG cell showing lack of an effect of DL-TBOA (averages of 18-26 events). The PG cell in this recording, the same as that shown in **Fig. 1D**, displayed bursts of EPSCs following OSN stimulation (**Fig. 1D2**), implying that the analyzed EPSCs reflected eTC inputs. (**C**) Summary of DL-TBOA-effects on eTC-driven currents in MC/eTCs and PG cells. MC/eTC data were separated by when the spiking eTC displayed ≥ 2 spikes versus 1 spike. \**p* = 0.0013 in paired t-test, comparing before and during DL-TBOA application, Bonferroni correction for multiple comparisons, *n* = 5 pairs. (**D**) Example eTC-MC pair recording showing reversible reduction of *EC_extra_* after sequential addition of the AMPA and NMDA receptor antagonists (NBQX, 10 M, and DL-AP5, 50 M). Note that the drugs reduced *EC_extra_* without reducing the number of spikes in the eTC. eTC spiking was induced by a direct depolarizing current step during a whole-cell recording. (**E**) Pharmacological analysis of light-evoked MC currents in cholecystokinin (CCK)-Cre mice. (**E1**) Confocal analysis of an olfactory bulb section taken from a CCK-Cre-tdTomato mouse. Note that tdTomato expression appears mainly in cells in the outer portion of the EPL (likely superficial tufted cells) and in the glomerular layer (likely eTCs). (**E2**) Sequential addition of DL-AP5 and NBQX reduced light-evoked (100 sec, 473 nm LED) excitatory currents in a MC from a CCK-Cre-ChR2 mouse. (**F**) The combined addition of NBQX and DL-AP5 eliminated ∼70% of *EC_extra_* regardless of the magnitude of the control current (linear regression fit: *r* = 0.08, *n* = 17, *p* = 0.77). Open circles = MC currents in eTC-MC pairs; open triangles = light-evoked currents in MCs; closed squares = eTC currents in eTC-eTC pairs. Each data point reflects one experiment. (**G**) Summary of GluR blocker effects on *EC_extra_* (all experiment-types combined). Application of NBQX (*left*; \**p* < 0.01 in Wilcoxon matched-pairs signed-rank test, *n* = 11) or DL-AP5 (*right*; *p* < 0.05 in Wilcoxon matched-pairs signed-rank test, *n* = 6) alone reduced *EC_extra_* by 50-70%, while the combination of NBQX+DL-AP5 generally elicited only modest additional decreases. Asterisks for the NBQX+DL-AP5 condition denote *p* < 0.05 with respect to control. (**H**) When only the eTC-MC and eTC-eTC pair recordings were analyzed (*n* = 9), clear effects of NBQX and NBQX+DL-AP5 were observed on *EC_extra_* (*left*), while spiking in the eTC was unaffected (*right*). \**p* < 0.02, Wilcoxon matched-pairs signed-rank test, *n* = 7.

In addition to glutamate accumulation, another potential contributor to the supralinear increase in *EC_extra_* are the properties of the extrasynaptic glutamate receptors. NMDA receptors in particular have well-established non-linear properties, owing to their voltage-dependent blockade by magnesium (Mayer et al., 1984; Nowak et al., 1984). NMDA receptors are also often located extrasynaptically (Clark et al., 1997; Kullman and Asztely, 1998) where, because of their comparatively high affinity for glutamate, they can respond to low micromolar concentrations of glutamate. To examine the contribution of different glutamate receptor conductances to *EC_extra_*, antagonists of AMPA (NBQX, 10-20 μM) and NMDA (DL-AP5, 50-100 μM) receptors were applied in different sequences during whole-cell recordings in eTC-MC and eTC-eTC pairs (*n* = 9; example in **Fig. 6D**; *V_hold_* = –77 mV). To increase throughput in this pharmacological analysis, some additional measurements combined single MC recordings with optogenetic stimulation of eTCs and other TCs in CCK-Cre-channelrhodopsin-2 (ChR2) transgenic mice (*n* = 8; **Fig. 6E1,E2**). These experiments took advantage of the expression of the neuropeptide cholecystokinin (CCK) in tufted cells (Liu and Shipley, 1994; see *Methods* and **Supplementary Fig. 2**). Across the pair-cell and optogenetic recordings, the combination of NBQX and DL-AP5 caused reversible reductions in most (∼70%) of the evoked current (**Fig. 6D,E2,F**) with no clear differences in drug effects for different types of stimulation methods or recording cell type (**Fig. 6F**). These effects on *EC_extra_* occurred without changes in the number of spikes in the spiking eTC (1±2% decrease in spikes due to NBQX + DL-AP5 in 9 pair-cell recordings; **Fig. 6D,H**). Importantly, we found that the effects of NBQX and DL-AP5 were not additive. While both NBQX and DL-AP5 when applied alone caused large current reductions (**Fig. 6D,E2,G**; 53 ± 7% due to NBQX, *n* = 11; 64 ± 7% due to DL-AP5, *n* = 6; *p* < 0.05 in Wilcoxon matched-pairs signed-rank test for both), the addition of the second drug caused only modest further decreases (16 ± 5% reduction of the total current when DL-AP5 was applied after NBQX, *n* = 11; 8 ± 3% reduction of the total current when NBQX was applied after DL-AP5, *n* = 5; **Fig. 6G**). Absence of additivity would be expected if *EC_extra_* mainly reflected ion flow through NMDA receptors and a major role of AMPA receptors was to mediate depolarization-induced unblocking of magnesium from NMDA receptors. In experiments in which DL-AP5 was applied first, the drug sometimes caused the preferential elimination of a slow current component (in 3 of 6 experiments, including example in **Fig. 6E2**), consistent with the different kinetics of NMDA versus AMPA receptors.

We considered one additional, common test of whether responses are mediated by NMDA receptors, which is to record currents at depolarized holding potentials. This manipulation can enhance NMDA receptor-mediated currents by relieving blockade of magnesium (Forsythe and Westbrook, 1988). Unfortunately, these experiments were not very informative with respect to the role of NMDA receptors. Recording *EC_extra_* in MCs or eTCs at voltages ranging between –27 mV and +43 mV had virtually no effect on its magnitude, polarity, or duration (**Supplementary Fig. 3A,B1,C,D**). We attribute the lack of an effect of depolarization to the fact that much of *EC_extra_* reflected glutamate receptors on MCs/eTCs that were electrically coupled to the test MCs/eTCs (Schoppa and Westbrook, 2002; Christie et al., 2005; Hayar et al., 2005; Pimentel and Margrie, 2008; Gire et al., 2012); thus changing the membrane potential of the test cell would have little effect on the driving force for current flow through glutamate receptors (Schoppa and Westbrook, 2002). Depolarizing the membrane potential (to –27 mV) did cause the expected large change in the magnitude of spontaneous OSN-EPSCs in eTCs (reduction by 73 ± 9%; *p* = 0.0025 in paired t-test comparing effect of depolarization on OSN-EPSC versus *EC_extra_* in 4 eTC-eTC pairs; **Supplementary Fig. 3B2,C**).

The pharmacological analyses suggested that much of the supralinear increase in *EC_extra_* with eTC spike number could have reflected the biophysical properties of extrasynaptic NMDA receptors as well as glutamate dynamics in the extrasynaptic space. We also obtained evidence against some other plausible mechanisms. For example, increasing spike number in eTCs could have recruited new, transmitter-activated conductances; however, arguing against this scenario was the fact that the combined addition of NBQX and DL-AP5 reduced most (∼70%) of the MC current for all-sized currents (**Fig. 6F**). The non-linear increase in *EC_extra_* also appeared to be largely independent of recurrent excitation involving secondary activation of other glutamatergic cells at the glomerulus. Because at least a major portion of eTC-to-MC transmission is due to spillover at eTC-to-PG cell synapses (see above), the recruitment mechanism predicted that, during recordings in eTC-PG cell pairs, the PG cell should have displayed rapid EPSCs not directly associated with spikes in the test eTC. Such events, reflecting spike activity in other eTCs excited by glutamate release from the test eTC, were only infrequently observed (when network-wide LLD events were excluded in the analysis; **Fig. 7**). Using derivative methods to detect rapid EPSCs during eTC spike bursts (with ≥ 2 spikes), we found that the vast majority of EPSCs in the PG cell occurred in a window 0.5-3.5 ms after each of the spikes (**Fig. 7A1,B1**), consistent with monosynaptic transmission from the test eTC. Analyzing the results on a burst-wide basis, most spike bursts in eTCs (77 ± 6%, *n* = 5 eTC-PG cell pairs) had no associated EPSCs in PG cells outside of the 0.5-3.5 ms window (**Fig. 7C**), and those bursts that had extra events generally had only 1 or 2. Overall, EPSCs outside of the 0.5-3.5 ms post-spike window occurred at a frequency (7.2 ± 2.6 Hz; *n* = 5) that was at most only modestly higher than the background EPSC rate in shuffled trials (2.0 ± 1.3 Hz; *p* = 0.07 in paired t-test). These results indicated that most of the supralinear rise in the extrasynaptic currents that we studied did not result from recurrent excitation (see *Discussion*).

**Figure 7.**
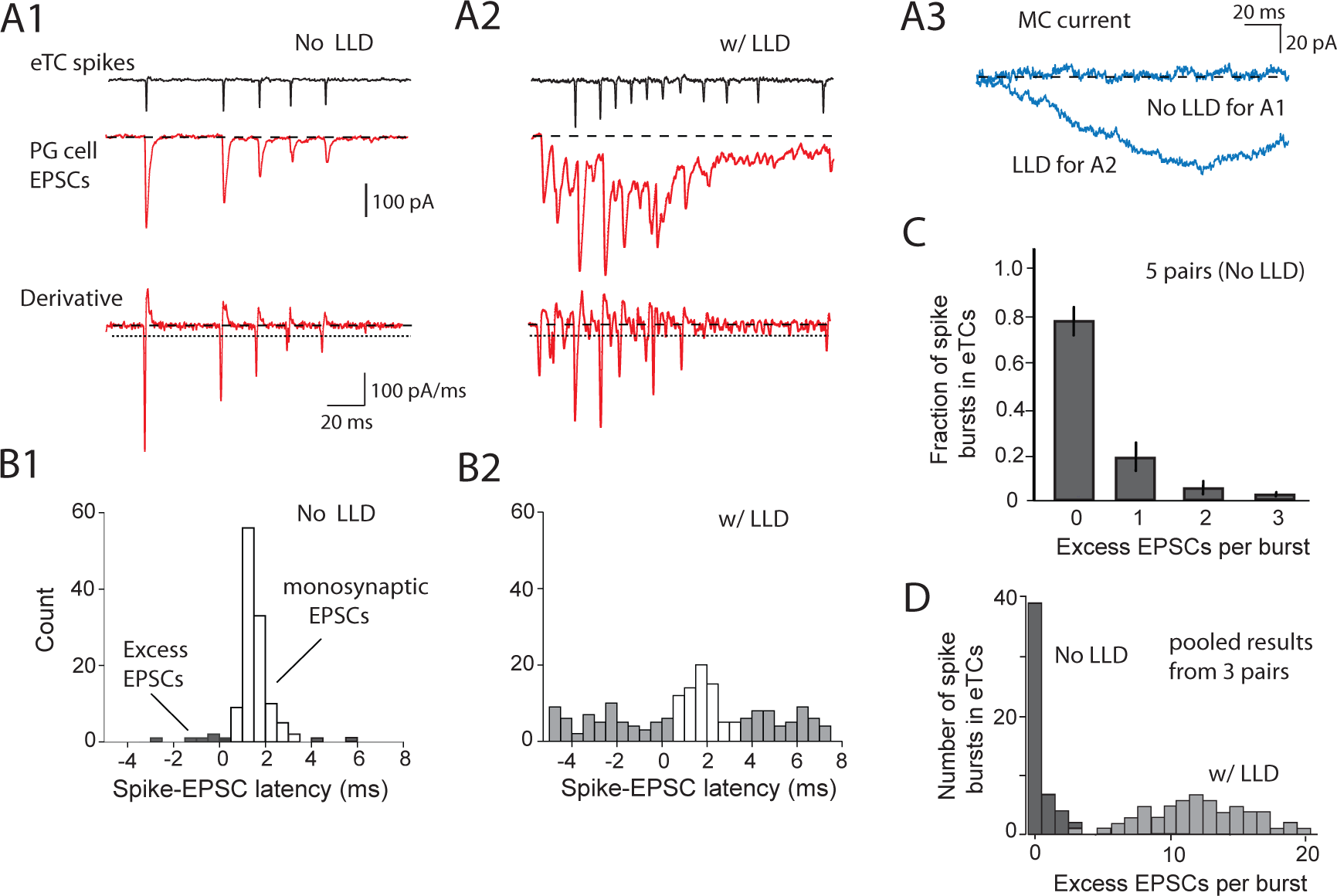
Supralinear increase in *EC_extra_* was mainly not due to recurrent excitation. (**A**) Examples of spontaneous currents from an eTC-PG cell-MC triple cell recording used to evaluate recurrent excitation. The eTC and PG cell bursts in **A1** and **A2** are differentiated by whether there was a co-occurring LLD in the glomerulus, as assessed from the MC current (**A3**). Note that in the burst in **A1** without an LLD, the PG cell appears to display no EPSCs besides those locked to spikes in the test eTC. The absence of excess EPSCs, consistent with no recurrent excitation, can also seen in the PG current derivative trace at bottom that was used for EPSC detection (based on threshold crossings; see horizontal dotted line). The burst in **A2** that was associated with an LLD had many excess EPSCs, but our analysis of *EC_extra_* excluded LLDs (see main text). (**B**) Histograms reflecting latencies between eTC spikes and PG cell EPSCs (peaks of derivative) for the experiment in **A**. Latencies of 0.5-3.5 ms (open bars) were considered to reflect monosynaptic transmission from the test eTC to the PG cell. Latencies derived from eTC-PG cell bursts that were both not associated with LLDs (**B1**) and associated with LLDs (**B2**) are illustrated. (**C**) Distribution of the number of PG cell EPSCs per burst that were not locked to eTC spikes across 5 eTC-PG cell pair recordings. Note that the large majority of burst-events that were not associated with LLDs also did not display excess PG cell EPSCs reflecting recurrent excitation. (**D**) In 3 dual or triple-cell recordings in which direct information was available about the LLD (pooled data), bursts associated with LLDs consistently had numerous excess EPSCs in the PG cell.

A potential caveat in using the eTC-to-PG cell pair recordings as a test for recurrent excitation was that the analysis assumed that each PG cell is well-connected to the population of eTCs at its target glomerulus and thus capable of reporting glutamate release from other cells. Supporting this assumption were responses in eTC-PG cell pairs when their target glomerulus engaged in LLDs, which, as discussed above, involve concerted excitation of all MCs and eTCs at a glomerulus. The LLDs were readily apparent as spontaneous events in three of the experiments, including the two MC-PG cell-eTC triple-cell recordings in which the MC current could be used to report the LLD (**Fig. 7A3**). LLDs were without exception associated with complex event clusters with numerous PG cell EPSCs uncoupled to spikes in the test eTC (**Fig. 7A2,B2,D**). The large number of uncoupled EPSCs in PG cells that co-occurred with LLDs indicated that the test PG cells could in fact report activity in other eTCs.

### Excitation and inhibitory responses to OSN stimulation

As a last step in our analysis, we tested whether the dynamic changes in excitation of different cell-types as a function of eTC spike number observed in the pair-cell recordings (summarized in **Fig. 5F**) were associated with corresponding changes in glomerular extrasynaptic excitation and inhibition (the “E/I balance”) in response to stimulation of OSN axons. The prediction was that low levels of OSN input, which presumably would drive few eTC spikes (see below), would lead to greater PG cell-mediated inhibition than extrasynaptic excitation but the balance would shift toward extrasynaptic excitation with greater OSN input. As a readout for extrasynaptic excitation and inhibition in a glomerulus driven by OSN input, we used voltage-clamp recordings in single eTCs (**Fig. 8A,B**). Recordings at hyperpolarized holding potentials (–77 mV) in these cells provided measures of both *EC_extra_*, likely originating from other eTCs (**Fig. 4F**), as well as an evoked OSN-EPSC that enabled us to estimate the co-occurring level of OSN input. At depolarized voltages (+28 mV), we could isolate an inhibitory current (*IC*) that reflected polysynaptic inhibition from local GABAergic cells (Zak et al., 2015; see *Methods* and **Supplementary Fig. 4**). *IC* likely reflected PG cell-to-eTC inhibitory synapses (Murphy et al., 2005), given that eTCs lack lateral dendrites and do not receive inputs from GABAergic granule cells. In addition, in parallel experiments conducted in GAD65-Cre-NpHR3 mice in which GAD65-positive PG cells (Kiyokage et al., 2010) expressed the inhibitory opsin halorhodopsin (NpHR3), light-pulses applied to the glomerulus suppressed most of *IC* (by 70 ± 8%; *p* = 0.004 in paired t-test, *n* = 4). In contrast to eTC currents, the current responses of MCs and other tufted cells are much more difficult to interpret in the context of the glomerular microcircuit. These other cells, which have lateral dendrites, receive granule cell inputs; also, MCs do not reliably report OSN activity driven by weaker stimuli (Najac et al., 2011; Gire et al., 2012; Vaaga and Westbrook, 2016). For all experiments, weak intensities of electrical stimulation were used (1 – 50 μA) and test eTCs were chosen that were associated with glomeruli at the surface of the slice. This enabled us to stimulate OSNs at one glomerulus with little or no activity in neighboring glomeruli (McGann et al., 2005; Gire and Schoppa, 2009; see *Methods* and **Supplementary Fig. 5**).

**Figure 8.**
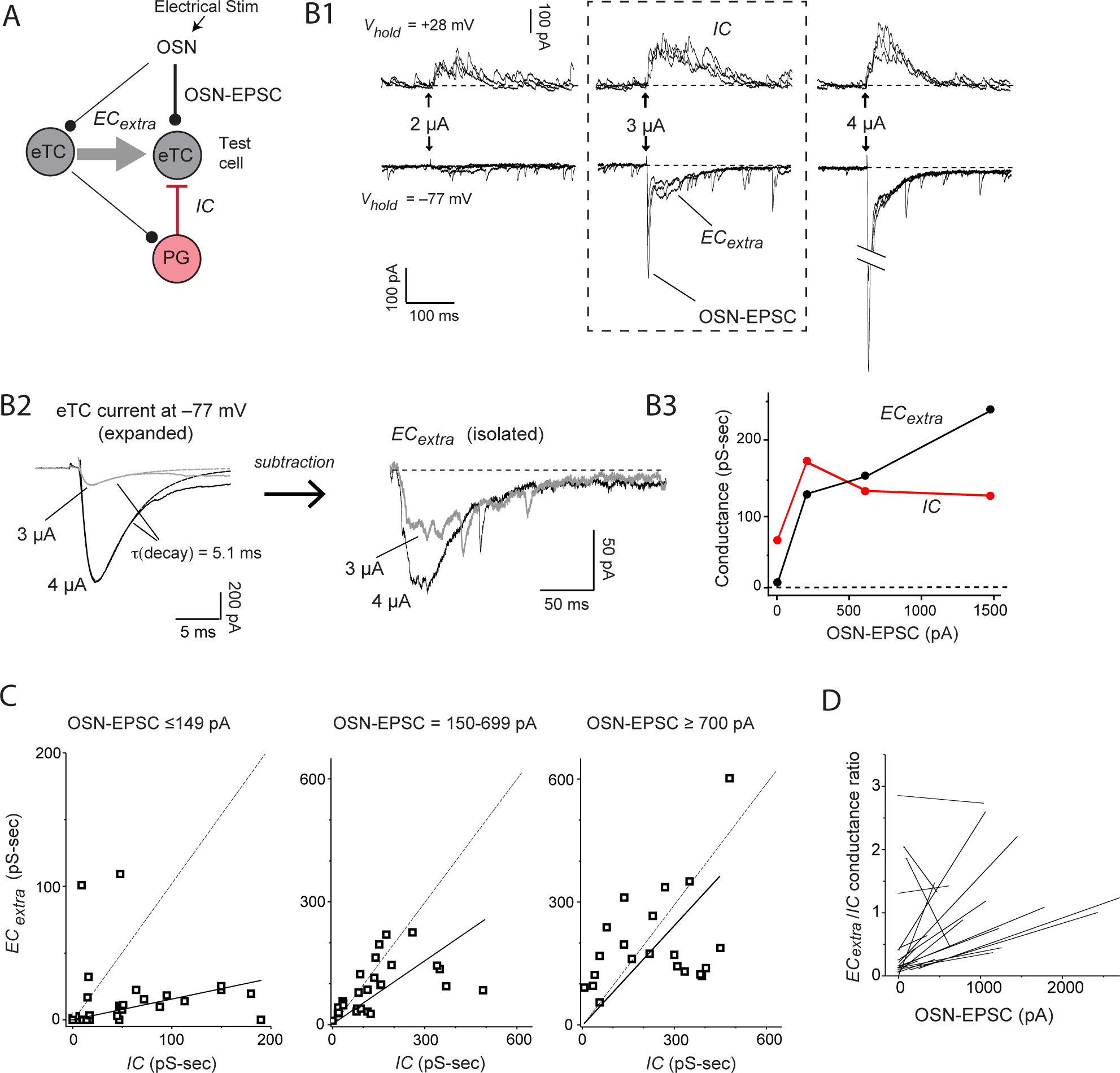
*EC_extra_* and synaptic inhibition following OSN stimulation. (**A**) Simplified circuit reflecting currents measured in a voltage-clamped eTC following stimulation of OSNs. Components include the monosynaptic OSN-EPSC, *EC_extra_* due to activation of other eTCs (OSN-to-eTC-to-eTC), and a polysynaptic inhibitory current (*IC*) due mainly to feedback inhibition from PG cells (OSN-to-eTC-to-PG cell-to-eTC). An early component of the inhibition could have reflected a feedforward pathway (not shown) mediated by ∼30% of PG cells directly excited by OSNs (Shao et al., 2009). (**B**) Example eTC current responses. (**B1**) Overlaid trials (3 each) of excitatory currents (at *Vhold* = –77 mV) and inhibitory currents (at *Vhold* = +28 mV) evoked by OSN stimulation at three intensities (2, 3, and 4 A). The 3-A data in boxed region best illustrate the current components defined in part **A**. Note that at the lowest intensity (2 A), inhibition was much larger than excitation and the OSN-EPSC was barely detectable. (**B2**) *EC_extra_* (*right*) was isolated by fitting the rise and most of the decay of the OSN-EPSC seen the composite current (*left*) with a sum of two exponentials, then subtracting the derived estimate of the OSN-EPSC. Average currents (5 trials) at two stimulus intensities are illustrated. (**B3**) Integrated conductances associated with *EC_extra_* and *IC* as a function of OSN-EPSC amplitude for this experiment. Each point reflects mean of 5 trials. (**C**) Summary of *EC_extra_* and *IC* conductance measurements across 21 eTC recordings, sorted by amplitude of the co-occurring OSN-EPSC. Solid lines reflect linear regression fits of data points (one per recording), constrained to pass through the origin, added here for illustrative purposes. Dashed lines reflect unity. (**D**) Linear regression fits of OSNEPSC amplitude versus *EC_extra_*/*IC* conductance ratio measurements from 19 individual eTC recordings. Each eTC recording had at least three OSN stimulation intensities sampled (actual data points not shown). Note positive slopes in all but 3 experiments, indicating that the *EC_extra_*/*IC* conductance ratio generally rose with increasing OSN input.

Relating *EC_extra_* and *IC* to the OSN-EPSC in eTCs revealed profound changes in the E/I balance as a function of OSN activity. Electrical stimuli that generated small OSN-EPSCs (peak absolute amplitudes ≤ 149 pA) resulted in responses that were nearly always dominated by inhibition (median *EC_extra_*/*IC* conductance ratio = 0.14, mean ± SEM = 0.53 ± 0.30, 30 OSN-EPSC values pooled from 19 eTC recordings; *p* < 0.01 in Wilcoxon matched-pairs signed-rank test in comparison to unity; **Fig. 8B1,B3,C**). However, larger OSN-EPSCs (150-699 pA) were associated with the emergence of large *EC_extra_* responses and *EC_extra_*/*IC* conductance ratios that were approaching unity (median ratio = 0.76, mean ± SEM = 0.67 ± 0.08, *n* = 27). Similar results were observed for the largest OSN-EPSCs (> 700 pA; median ratio = 0.99, mean ± SE = 1.32 ± 0.44, *n* = 21; *p* > 0.10 in Wilcoxon matched-pairs signed-rank test for both medium and large OSN-EPSCs). The shift in the *EC_extra_*/*IC* conductance ratios could also be observed on an experiment-wide basis. In 19 eTC recordings in which at least 3 OSN activity levels were sampled (≥ 3 OSN stimulus intensities), the conductance ratios nearly always rose as a function of OSN activity (slope of fitted lines = 3.5/nA; mean ± SE = 3.6 ± 2.0/nA, *n* = 19; *p* < 0.01 in Wilcoxon matched-pairs signed-rank test in comparison to zero slope; **Fig. 8D**).

The *EC_extra_*/*IC* ratios following OSN stimulation appeared to match the predictions based on the excitatory current recordings in eTC-MC/eTC and eTC-PG cell pairs (compare plots in **Fig. 8B3,D** and **Fig. 5F**), but were the observed changes in fact related to each other? This was tested, first, by verifying the assumption that the measures of stimulus strength in the two sets of experiments – the amplitude of the OSN-EPSC and the number of eTC spikes – were correlated. Using whole cell recordings in eTCs that examined responses to OSN stimulation while switching between voltage-clamp and current-clamp, we indeed found this to be the case. Increasing OSN-EPSCs were associated with both a higher probability of eTC spiking (**Supplementary Fig. 6A**) and a greater number of eTC spikes when spikes were evoked (mean correlation coefficient comparing OSN-EPSC and evoked number of spikes, excluding failures = 0.62, *n* = 16 eTCs, *p* = 0.0032; **Supplementary Fig. 6B**). Furthermore, *EC_extra_* and *IC* evoked in eTCs by OSN stimulation displayed a similar sensitivity to the glutamate uptake inhibitor DL-TBOA as the excitatory currents recorded in the eTC-MC/eTC and eTC-PG cell pairs (**Fig. 6A-C**). While normalizing for the strength of OSN input, DL-TBOA (10 μM) caused large increases in *EC_extra_* evoked by OSN stimulation (**Fig. 9A,B1;** 3.2-fold increase in slope of lines fitting OSN-EPSC versus *EC_extra_* values; *p* < 0.001 in ANCOVA, *n* = 26 for control, *n* = 21 for TBOA; pooled results from six eTC recordings) but did not impact *IC* (**Fig. 9B2**; *p* > 0.10 from ANCOVA). DL-TBOA also caused large increases in the *EC_extra_*/*IC* conductance ratios from the same experiments (**Fig. 9B3,C**; *p* < 0.001 in ANCOVA). These results suggest that the dynamic features associated with extrasynaptic excitation that contributed to the shifts in excitation of excitatory and inhibitory cells in the pair-cell recordings also contributed to changing the E/I balance within a glomerulus in response to different levels of OSN input.

**Figure 9.**
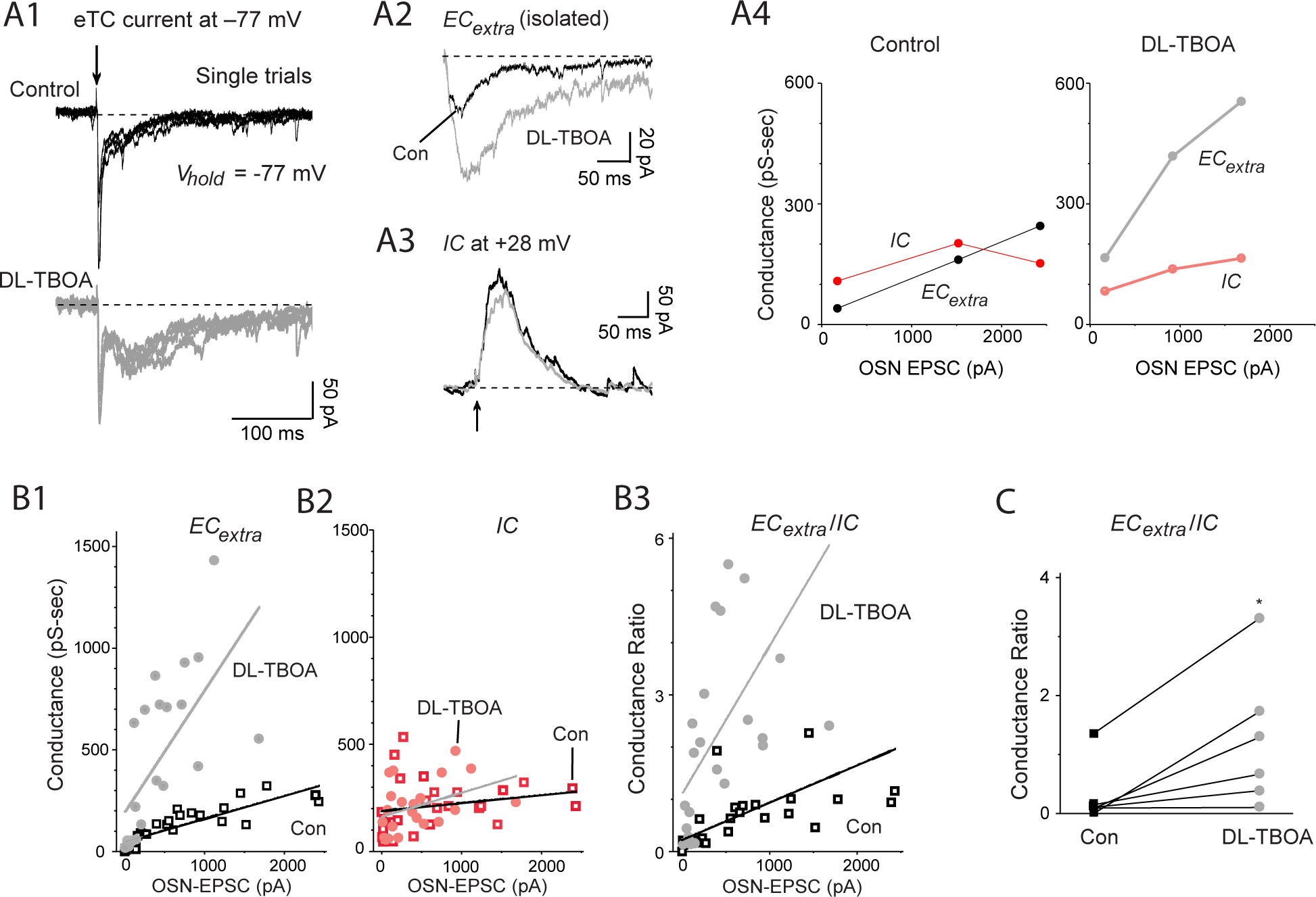
Sensitivity of eTC currents evoked by OSN stimulation to glutamate uptake blockade. (**A**) Effect of DL-TBOA in example eTC current recording. (**A1**) Excitatory currents in an eTC evoked by OSN stimulation (2.5 μA; superimposed single trials) recorded under control conditions (top) and following bath application of DL-TBOA (10 μM; bottom). (**A2** and **A3**) DL-TBOA enhanced *EC_extra_* (average of 5 trials; isolated as in **Fig. 8B2**) but not the inhibitory current *IC* recorded at +28 mV. (**A4**) Summary of the effects of DL-TBOA in this experiment. Note the especially marked change in the balance between *EC_extra_* and *IC* due to DL-TBOA when the OSN-EPSC was the smallest. (**B**) Summary of DL-TBOA effect across six eTC recordings. Plotted are the OSNEPSC amplitudes versus the integrated conductances associated with *EC_extra_* (**B1**) and *IC* (**B2**), as well as the *EC_extra_/IC* conductance ratio (**B3**). Linear regression fits of the data points (overlaid) indicated that DL-TBOA enhanced *EC_extra_* and the *EC_extra_/IC* conductance ratio (*p* < 0.001 from ANCOVA for both) but not *IC* (*p* > 0.10). There are more data points in each plot than the number of recordings reflecting the fact that more than one OSN stimulus intensity was sampled in most recordings. (**C**) *EC_extra_/IC* conductance ratios separated by experiment under control conditions and in DL-TBOA. For experiments in which more than one OSN stimulus intensity was sampled, the indicated ratios reflect averages of the ratios at each stimulus intensity. \**p* < 0.05 in Wilcoxon matched-pair signed-rank test, *n* = 6.

## Discussion

Recent evidence indicates that a major mechanism of sensory input-driven activation of MCs is an indirect pathway mediated by excitatory eTCs at glomeruli (De Saint Jan et al., 2009; Najac et al., 2011; Gire et al., 2012). Further, the eTC-to-MC transmission step in the pathway occurs not via conventional synapses, but rather extrasynaptically. Here we have studied the underlying mechanisms of extrasynaptic transmission in bulb slices, as well as its dynamics with respect to inhibition in glomeruli with changing levels of eTC activity and sensory input. Our results point to a local intraglomerular thresholding mechanism that could have a number of important functions during natural odor-evoked responses.

### Mechanisms and dynamics of extrasynaptic excitation

We found that at least a major component of extrasynaptic eTC-to-MC transmission reflects glutamate released at dendrodendritic eTC-to-PG cell synapses acting on extrasynaptic receptors on MC apical dendrites. This conclusion was based, first, on pair- and triple-cell patch-clamp recordings in which slow excitatory currents were observed in MCs that were time-locked and correlated in amplitude to fast EPSCs in PG cells. Second, MCs displayed small currents time-locked to mEPSCs in PG cells recorded in TTx. Because mEPSCs reflect glutamate release at single release sites, the time-locked currents in MCs must have reflected spillover. Third, we found evidence for an ultrastructual correlate of spillover. In studies with labeled eTC dendrites, complexes within glomeruli were observed that included an eTC synapse onto a GABAergic dendrite (likely that of PG cells) and a presumed MC dendrite in close proximity. Notably absent from the complexes were astroglial processes separating the MC dendrite from the eTC-to-PG cell synapse that function to limit spillover of glutamate through activity of their transporters (Asztely et al., 1997; Murphy-Royal et al., 2017). Instead, glial processes were observed to sometimes surround the eTC-PG cell-MC dendritic complexes.

A potential concern with our experiments, conducted in brain slices, is that the prominence of extrasynaptic signaling was an artifact of the brain slice preparation. In hippocampus, it has been reported that certain types of brain slicing procedures can cause glial processes to retract (Bourne and Harris, 2011); such an effect might promote extrasynaptic signaling. In this light, our ultrastructural results, which were based on eTC dye-fills in rat olfactory bulb slices prepared in the same manner as those used in the physiology experiments, are particular noteworthy since they showed that glial processes were present. In addition, other ultrastructural studies conducted using conventional whole-brain fixation methods (Chao et al., 1997; Kasowski et al., 1999) have reported glial-wrapped subcompartments in glomeruli resembling ours that included unlabeled glutamatergic and GABAergic dendrites. A last point is that we found in our physiological studies that spillover was not a general phenomenon in our brain slices. In eTC-MC pair-recordings, there was no evidence for spillover-mediated responses in MCs from glutamate released at OSN-to-eTC synapses.

As a means of signaling between neurons, glutamatergic activation of extrasynaptic AMPA and NMDA receptors has been reported in a wide range of circuits (Asztely et al., 1997; Isaacson, 1999; Carter and Regher, 2000; DiGregorio et al., 2002; DeVries et al., 2006; Szapiro and Barbour, 2007; Chalifoux and Carter, 2011; Coddington et al., 2013; Wild et al., 2015). The extrasynaptic, spillover-mediated transmission that we have described between eTCs and MCs however stands out, first, because it is the *primary* mode of signaling between eTCs and MCs. Extrasynaptic transmission in most systems occurs as a means to amplify co-occurring direct synaptic transmission, but eTCs form very few if any direct synaptic connections onto MCs (Pinching and Powell, 1971; Bourne and Schoppa, 2017). To our knowledge, only two other examples of glutamatergic transmission dominated by extrasynaptic mechanisms have been described, from climbing fibers to molecular layer interneurons in the cerebellum (Szapiro and Barbour, 2007) and between MCs in the bulb (Isaacson, 1999), and, in the latter case, recordings were conducted in zero extracellular magnesium which favored extrasynaptic NMDA receptor-mediated signaling (the study also did not explicitly test for spillover). The eTC-to-MC extrasynaptic signaling that we characterized also differs from examples in other circuits in that it is part of a major pathway of information flow (OSN-to-eTC-to-MC) and also because of the unusual synaptic arrangement. Spillover-mediated transmission between two types of excitatory cells (from eTCs onto MCs) occurs as a result of glutamate released at synapses onto *inhibitory* cells (GABAergic PG cells). Functionally, such an arrangement means that the system is set up so that any episode of MC excitation co-occurs with at least some level of excitation of inhibitory cells in close proximity.

In addition to providing information about the basic mechanisms of extrasynaptic signaling between eTCs and MCs, our recordings also provided quantitative information about the dynamics of extrasynaptic excitation with changing eTC spike number. As expected from the long distance between glutamate release sites and extrasynaptic glutamate receptors, *EC_extra_* elicited by single eTC spikes was very small but rose in a highly supralinear fashion with increasing spikes. Mechanistically, this supralinearity in the rise of *EC_extra_* appeared to reflect at least in part the dynamics of glutamate accumulation at sites of extrasynaptic glutamate receptors (Asztely et al., 1997; Carter and Regehr, 2000; Clark and Cull-Candy, 2002; Pendyam et al., 2009). Based on our finding that NMDA receptors contributed to most of *EC_extra_*, it is likely that the non-linear biophysical properties of these receptors (Mayer et al., 1984; Nowak et al., 1984) also contributed to the supralinearities. An interesting – perhaps surprising – feature of the supralinear rise in *EC_extra_* with eTC spike number was that it occurred in spite of the fact that each successive eTC spike appeared to produce smaller glutamate transients. If the EPSC in the PG cell is taken to reflect the relative amount of glutamate release, our results suggest that each successive eTC spike generated 15-30% of the amount of glutamate released by the first spike (the values for the second and fourth spikes, respectively, were 31 ± 2% and 17 ± 1%, *n* = 5). We suggest that this result may be explained if the mechanisms underlying the supralinearities (glutamate accumulation and NMDA receptor biophysics) are just exceptionally strong such that even small additional increases in glutamate caused large increases in *EC_extra_*. It is also possible that there are mechanisms yet to be identified that increase the potency of eTC spike bursts. For example, negative coupling between neural activity during spike bursts and glutamate uptake by astroglial transporters has been reported (Armbruster et al., 2016). Another scenario that we cannot completely exclude is one in which repetitive spiking in eTCs recruits the release of excitatory neurotransmitters other than glutamate (Ma et al., 2013). In our experiments, we found that blockade of ionotropic glutamate receptors reduced most (∼70%) of even the largest *EC_extra_* responses (**Fig. 6F**), which was inconsistent with a significant role for other neurotransmitters. Also, the glutamate uptake inhibitor DL-TBOA prolonged the larger *EC_extra_* responses (**Fig. 6A,C**). However, it is possible that other neurotransmitters work synergistically with glutamate to enhance *EC_extra_*.

In many neural circuits, a major contributor to supralinear increases in excitation are recurrent mechanisms in which glutamate release from one neuron drives spike activity in other excitatory neurons (McCormick et al., 2015; Dehghani et al., 2016). Recurrent excitation did not appear to play a major role in driving the supralinearities that we studied here. Using the PG cell current response during eTC-PG cell recordings as a reporter of activity amongst the network of excitatory cells at a glomerulus, spike bursts in the test eTC were typically not associated with activity in other excitatory cells, as would be required for a recurrent mechanism. At the same time, it should be emphasized that recurrent excitation is widely recognized to be an important component of the MC excitatory response to OSN input. MCs and eTCs engage in large, all-or-none LLDs that co-occur among all excitatory cells at a glomerulus and that reflect recurrent excitation (Carlson et al., 2000; Schoppa and Westbrook, 2001; Gire and Schoppa, 2009). We would suggest that the extrasynaptic excitation that we studied here fits into this broader picture of glomerular excitation by providing the “seed” that triggers the LLD. Evidence for such a triggering role was occasionally observed when *EC_extra_* temporally preceded the much larger LLD (**Fig. 4A3**).

### Intraglomerular E/I balance and information processing

In order to understand the potential function of extrasynaptic excitation in glomeruli, we compared *EC_extra_* to the activity of GABAergic PG cells in relation to the strength of a stimulus. This was done in two ways, first in a series of pair-cell recordings in which we related the magnitude of the excitatory currents in MC/eTCs versus PG cells to the number of spikes in an eTC. This provided a measure of the relative strength of extrasynaptic excitation versus the excitatory drive on inhibitory cells. In addition, we measured *EC_extra_* and PG cell-mediated inhibitory currents in single eTCs in response to different levels of OSN stimulation at one glomerulus. In these latter experiments, direct information about OSN activity was available in each recording from the size of the OSN-EPSC that co-occurred with *EC_extra_* and the inhibitory currents. Both types of experiments yielded the same basic result (**Fig. 5F** and **Fig. 8**): Weak stimuli a (low number of eTC spikes or small OSN-EPSCs) yielded an E/I balance that significantly favored inhibition, but stronger stimuli (high eTC spike number or large OSN-EPSCs) caused a significant shift in favor of extrasynaptic excitation of excitatory cells. Mechanistically, the shift in the E/I balance resulted from both the supralinear rise in *EC_extra_* (discussed above) together with presynaptic depression at eTC-to-PG cell synapses. It is possible that other mechanisms that we did not investigate here also contributed to the shifting E/I balance, for example, differences in cell input resistances (Cleland and Sethupathy, 2006).

From the point of view of odor coding, the shifting E/I balance within one glomerulus could have a number of important functions, for example contributing to the narrow tuning of output MCs and TCs to different odors (Yokoi et al., 1995; Davison and Katz, 2007; Tan et al., 2010). If eTC spike number and OSN-EPSC amplitude are taken as surrogates for the affinity of an odor to a specific OR, our results would imply that a low-affinity odor would mainly drive inhibition within an OR-specific glomerulus while a high-affinity odor would be much more likely to cause excitation. A single glomerulus could thereby act as a threshold that selectively enables output MCs/TCs to spike in response to a high-affinity odor. **Figure 10** illustrates a somewhat more specific version of this thresholding effect that takes into account prior evidence for a prominent multi-step OSN-to-eTC-to-MC pathway (De Saint Jan et al., 2009; Najac et al., 2011; Gire et al., 2012), together with new evidence provided here that eTCs send extrasynaptic excitatory signals to each other (**Fig. 4F**). In the framework of this model, the locus where the E/I balance is most critical for determining whether a specific odor excites MCs is within the network of eTCs that lies between OSNs and MCs. The shifting intraglomerular E/I balance could have other functions besides acting as a threshold for odors of differing affinity for an OR, for example limiting bulbar responses to low concentrations of odor (where it would act as a concentration threshold) and/or contributing to sparse neural responses (Rinberg et al., 2006; Koulakov and Rinberg, 2011).

**Figure 10.**
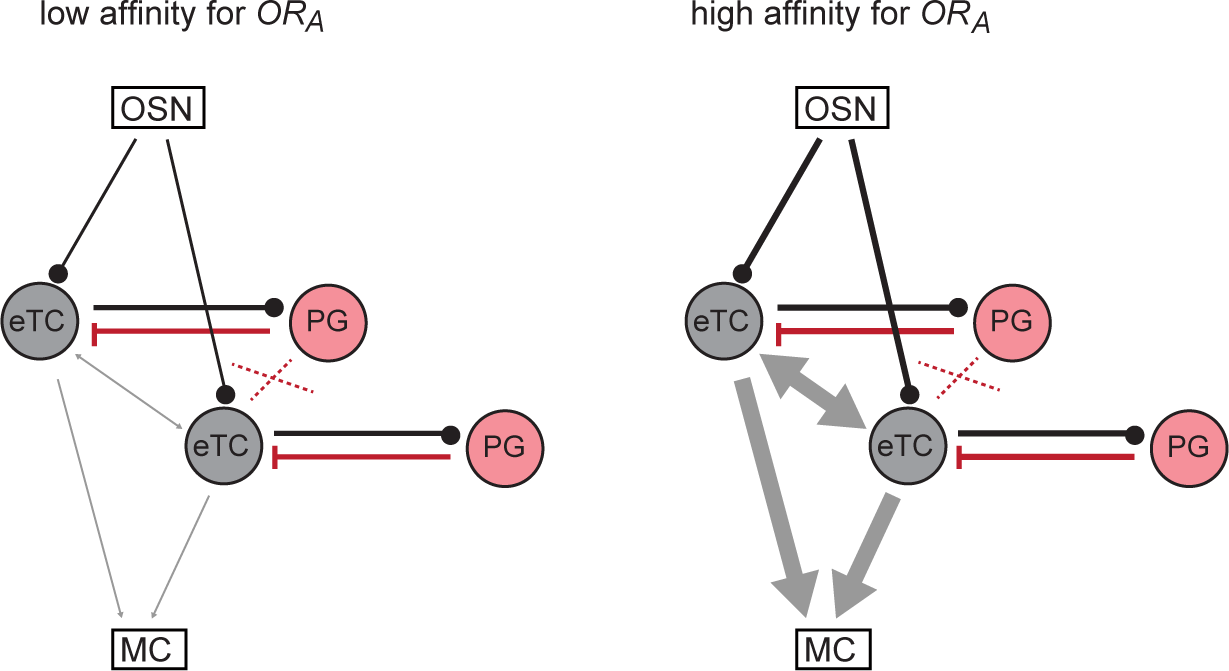
Conceptual model for narrowing the tuning MCs to odors of differing OR affinities. In the two simplified schemes, a glomerulus is shown that carries information about a particular OR (*OR_A_*). Signaling from OSNs to MCs occurs through a feedforward OSN-to-eTC-to-MC pathway. (*left*) An odor with a low affinity for *OR_A_* causes weak excitation of OSNs, in turn resulting in limited spiking in eTCs. This generates synaptic eTC-to-PG cell excitation and PG cell-to-eTC inhibition (black and red lines) that are much stronger than extrasynaptic eTC-to-eTC excitation (gray arrows). With limited recurrent excitation amongst eTCs, there is also little extrasynaptic excitation reaching MCs. (*right*) An odor with a high affinity for *OR_A_* excites eTCs more strongly, which causes only modest increases in the synaptic signals between eTCs and PG cells but supralinear increases in extrasynaptic eTC-to-eTC excitation. With substantial recurrent excitation amongst eTCs, a large extrasynaptic excitatory signal can also reach MCs.

As one considers the significance of the intraglomerular threshold in olfactory processing, a number of factors must be considered. First, a major premise of our proposed conceptual model needs to be emphasized, which is that information that originates in OSNs passes through eTCs prior to reaching MCs; it is then that the signals will be maximally impacted by the thresholding mechanism. A case can be made that this is not always the case, given that, in slice experiments, stimulation of OSNs can elicit spikes in MCs that reflect direct OSN synapses (Najac et al., 2011). Nevertheless, the available evidence from slice studies supports that the multistep OSN-to-eTC-to-MC is a major pathway for information flow, indeed accounting for most of the excitatory current and spikes in MCs induced by OSN stimulation (Najac et al., 2011; Gire et al., 2012; Vaaga and Westbrook, 2016). This prominence reflects several factors, including the fact that OSN inputs produce much larger direct monosynaptic signals to eTCs than MCs (Gire et al., 2012; Vaaga and Westbrook, 2016), the high intrinsic excitability of eTCs (Liu and Shipley, 2008), and the large size of the feedforward eTC-to-MC signal (Najac et al., 2011; Gire et al., 2012). In addition, modeling studies that have attempted to replicate *in vivo* odor-evoked responses have included a multi-step pathway of exciting MCs (Fukunaga et al., 2012; Carey et al., 2015) and, in one case, required that direct OSN inputs into MCs be weak (Fukunaga et al., 2012). Thus it is highly likely that an intraglomerular threshold involving OSN-to-eTC-to-MC signaling functions in a significant capacity during odor-evoked responses.

Another consideration is that the possible role of an intraglomerular threshold in narrowing the tuning of MCs/TCs runs up against a significant literature that has proposed a very different tuning mechanism, involving lateral inhibition between glomeruli mediated by GABAergic granule cells (GCs; Yokoi et al., 1995; Luo and Katz, 2001; Tan et al., 2010; Koulakov and Rinberg, 2011; Yu et al., 2014). Comparatively, how much existing evidence is there for intra- versus interglomerular mechanisms for controlling the tuning of MCs to odors? Unfortunately, it is extremely difficult to obtain direct information about the relative contribution of each using *in vivo* approaches. Because an inherent feature of any odor-evoked response is that many glomeruli are activated (Yang et al., 1998; Uchida et al., 2000; Rubin and Katz, 2001), neither intra- nor interglomerular odor-evoked mechanisms can be studied in isolation. Nevertheless, there is indirect supportive evidence for an intraglomerular mechanism for controlling the tuning MCs/TCs. For example, an increase in odor concentration, which is similar to increasing odor affinity for an OR, can result in a sign-inversion in the MC/TC response from suppression to excitation (Fukunaga et al., 2014; Economo et al., 2016). This mimics the shift from inhibition to excitation that we observed with increasing OSN input (**Fig. 8**). In addition, optogenetic silencing of PG cells but not GCs can reduce odor-evoked suppression of MCs/TCs in anesthetized mice (Fukunaga et all, 2014). PG cells are axon-less, extend dendrites into only one glomerulus, and do not appear to be excited by interglomerular connections (Whitesell et al., 2013); hence their contribution to odor-evoked suppression (and thus odor tuning) must be mediated by strictly intraglomerular mechanisms. The efficacy of lateral inhibition in tuning MCs/TCs, at least of the center-surround form, can also be questioned based on the fact that glomeruli in the bulb are only loosely ordered topgraphically (Mori et al., 2006; Fantana et al., 2008; Soucy et al., 2009; Ma et al., 2012). Thus, on balance, there is arguably at least as much *in vivo* experimental evidence consistent with a role for an intraglomerular mechanism in controlling the tuning of MCs/TCs as there is for GC-mediated lateral inhibition (Abraham et al., 2010; Nunez-Parra et al., 2013; Economo et al., 2016). We would suggest that a possible scenario is that the two types of mechanisms work in concert. Because GCs are well excited by cortical feedback (Balu et al., 2007; Boyd et al., 2012), an intraglomerular mechanism might operate as the default to narrow MC/TC tuning, but GC-mediated lateral inhibition could further narrow MC/TC tuning under conditions of high cortical activity (Cazakoff et al., 2014; Otazu et al., 2015).

## Materials and methods

### Animals and slice preparation

Male and female 8-20 day-old Sprague Dawley rats obtained from Charles River Laboratories were used for most experiments. Optogenetic experiments were conducted in transgenic mice (20-27 day PN) that expressed ChR2 under the promoter for CCK or NpHR3 under the promoter for GAD65. These mice (called CCK-Cre-ChR2 and GAD65-Cre-NpHR3 in our study) were generating by breeding Ai32(RCL-ChR2(H134R)/EYFP) or Ai39(RCL-eNpHR3.0ChR2/EYFP) mice (Jackson Laboratories #012569 and #014539) with CCK-Cre or GAD65-Cre mice (Jackson Laboratories #012706 and #010802). CCK-Cre and GAD65-Cre mice were characterized anatomically by breeding these mice with floxed Ai14(RCL-tdTomato)-D mice (Jackson Labs, #007908), yielding CCK-Cre-tdTomato and GAD65-Cre-tdTomato mice. All experiments were conducted under protocols approved by the Animal Care and Use Committee of the University of Colorado, Anschutz Medical Campus.

Acute horizontal olfactory bulb slices (300-400 µm) were prepared following isoflurane anesthesia and decapitation. Olfactory bulbs were rapidly removed and placed in oxygenated (95% O_2_, 5% CO_2_) ice cold solution containing (in mM): 72 sucrose, 83 NaCl, 26 NaHCO_3_, 10 glucose, 1.25 NaH_2_PO_4_, 3.5 KCl, 3 MgCl_2_, 0.5 CaCl_2_ adjusted to 295 mOsm. Olfactory bulbs were separated into hemispheres with a razor blade and attached to a stage using adhesive glue applied to the ventral surface of the tissue. Slices were cut using a vibrating microslicer (Leica VT1000S) and were incubated in a holding chamber for 30 minutes at 32°C. Subsequently, the slices were stored at room temperature.

### Electrophysiological recordings

Experiments were carried out under an upright Zeiss Axioskop2 FS Plus microscope (Carl Zeiss MicroImaging) fitted with differential interference contrast (DIC) optics, video microscopy and a CCD camera (Hamamatsu). Identified cells were visualized with 10x or 40x Zeiss water-immersion objectives. Recordings were performed at 30-35°C.

The base extracellular recording solution contained the following (in mM): 125 NaCl, 25 NaHCO_3_, 1.25 NaHPO_4_, 25 glucose, 3 KCl, 1 MgCl_2_, 2CaCl_2_ (pH 7.3 and adjusted to 295 mOsm), and was oxygenated (95% O_2_, 5% CO_2_). The pipette solution for most whole-cell recordings contained: 125 K-gluconate, 2 MgCl_2_, 0.025 CaCl_2_, 1 EGTA, 2 Na_3_ATP, 0.5 Na_3_GTP, and 10 HEPES, (pH 7.3 with KOH and adjusted to 215 mOsm). For whole-cell recordings from eTCs, 30 mM glutamic acid was added to the pipette to prevent run-down of evoked glutamatergic currents (Ma and Lowe, 2007). For whole cell recordings of eTC responses to OSN stimulation, the K-gluconate in the pipette solution was replaced with an equimolar amount of cesium methanosulfonate, as well as the sodium channel blocker QX-314 (10 mM) to block action potentials. All whole-cell recordings included 100 µM Alexa 488 or, in a few cases, Alexa 594 in the pipette solution to allow for visualization of cell processes. Loose cell-attached recordings from eTCs were made with a pipette that contained the extracellular solution. Patch pipettes, fabricated from borosilicate glass, were pulled to a resistance of 3-4 MΩ for MCs, 4-6 MΩ for eTCs, and 6-8 MΩ for PG cells. Current and voltage signals in the single- and pair-cell experiments were recorded with a Multiclamp 700B amplifier (Molecular Devices, San Jose, CA), low-pass filtered at 1.8 kHz using an eight-pole Bessel filter, and digitized at 10 kHz. Triple-cell recordings also incorporated an Axopatch 200B amplifier (Molecular Devices). Data were acquired using Axograph X software on an Apple Mac Pro computer. All drugs were delivered via bath application at a flow rate constant to the baseline measurements.

Cell identity was determined in part by visualizing Alexa 488- or Alexa 594-mediated fluorescence signals. MCs were easily identified by their position in the MC layer and large cell bodies. eTCs were distinguished from other cells in the glomerular layer by their position in the inner half of the layer, their relatively large, spindle-shaped somata (≥ 10 µm diameter), a single highly branched apical dendrite and no lateral dendrite, and a relatively low input resistance (∼0.2 GΩ; Hayar et al., 2004b). PG cells were identified by their small soma (< 10 µm diameter), small dendritic arbor that was confined to one glomerulus, and high input resistance (> 0.5 GΩ; Hayar et al., 2004b; Murphy et al., 2005; Shao et al. 2009). In addition, we only considered PG cells that displayed bursts of EPSCs reflecting inputs from bursting eTCs (Hayar et al., 2004a; Shao et al., 2009; ∼70% of the total), either spontaneously or in response to OSN stimulation. Fluorescence measurements were performed under whole-field epi-illumination using a DG-4 light source (Sutter Instruments, Novato, CA). Signals were detected by a CoolSnap II HQ CCD camera (Photometrics, Tucson, AZ) under control of Slidebook (Intelligent Imaging Innovations, Denver, CO) software.

In multi-cell recordings, we determined that the cells sent their apical dendrites to the same glomerulus in part based on anatomical measurements (example images in **Figs. 1D3** and **2B2**). In the example image of the PG cell-MC pair in **Fig. 1D3**, both cells were filled with the same dye (Alexa-488); we were able to trace much of the small dendritic arbor of the PG cell by following its dendrites from the cell body. In other pairs, we generally did not determine the detailed anatomy of the apical dendrites of the different cells but simply verified that the cells sent their apical dendrites to the same glomerulus. For eTC-MC or eTC-eTC pairs, physiological evidence that the test cells were associated with the same glomerulus was also obtained from the presence of perfectly coincident LLDs (Carlson et al., 2000; Gire and Schoppa, 2009) that occurred either spontaneously and/or following OSN stimulation (see example in **Supplementary Fig. 1**). When an eTC was in LCA patch mode, LLDs in the MC perfectly co-occurred with long-lasting bursts of spikes in the eTC. For recordings from eTC-PG cell pairs affiliated with the same glomerulus, every burst of spikes in the eTC was associated with a burst of rapid EPSCs in the PG cell (Hayar et al., 2004a) and no bursts of EPSCs in the PG cell were observed without co-occurring bursts of spikes in the eTC.

For the optogenetic measurements of extrasynaptic currents in MCs in CCK-Cre-ChR2 mice (**Fig. 6E**), light pulses (0.1 ms; 473 nm LED) were applied through the 40x objective centered on the glomerulus targeted by the MC. Prior to these recordings, CCK-Cre mice were characterized from the fluorescence expression patterns observed in CCK-Cre-tdTomato mice under an Olympus Fluoview FV1000 confocal microscope. Td-Tomato expression appeared to be confined to different tufted cells (Liu and Shipley, 1994), especially eTCs in the glomerular layer and superficial tufted cells with cell bodies in the outer portion of the external plexiform layer (EPL; see **Fig. 6E1)**; middle tufted cells (mTCs) located in more inner regions of the EPL also displayed some td-Tomato expression. Cells in the mitral cell layer did not exhibit significant tdTomato expression in mice at ≥ PN 20 (the age of CCK-Cre-ChR2 mice used in the electrophysiological recordings). Selective expression of Cre in different subtypes of tufted cells was also verified using recordings of ChR2-mediated currents in CCK-Cre-ChR2 mice. Light pulses (20 ms-duration, 473 nm LED; applied through a 40x objective) elicited large inward currents in eTCs and mTCs (mean peak = –450 ± 105 pA, *n* = 10; **Supplementary Fig. 2**) with an onset time < 1 ms after the start of the light pulse. Such ChR2-mediated currents were never observed in MCs (*n* = 13).

For experiments employing OSN stimulation (**Figs. 1,8,9**), stimulation of OSN axons was performed using a broken-tip patch pipette (5-10 µm-diameter) placed in the olfactory nerve layer, 50-100 µm superficial to the glomerular layer. Current injections were delivered by a stimulus isolator (World Precision Instruments) under control of a TTL output from Axograph X software. In order to limit neural activity to the target glomerulus of the test eTC as much as possible, weak intensities of electrical stimulation (1-50 µA) were used, combined with choosing glomeruli for analysis that generally included clear, associated bundles of OSN axons (McGann et al., 2005; Gire and Schoppa, 2009). Excitatory and inhibitory currents were isolated, respectively, at holding potentials of –77 mV and +28 mV.

### Controls for OSN stimulation experiments

As control experiments supporting the analysis of eTC currents following OSN stimulation (**Figs. 8**,**9**), we first wanted to verify that the evoked outward currents at +28 mV were mediated mainly by PG cells. This was accomplished optogenetically in GAD65-Cre-NpHR3 mice, wherein we found that prolonged light pulses (590 nm LED, 320-ms pulses starting 20 ms prior to OSN stimulation applied in the glomerular layer) reduced inhibitory currents in eTCs (**Supplementary Fig. 4A**). Because PG cells preferentially express GAD65 (Kiyokage et al., 2010), the light-evoked suppression of inhibition in eTCs argued that at least a large component of the inhibition was mediated by PG cells. To verify that Cre was expressed in PG cells in the GAD65-Cre-NpHR3 mice, direct recordings were made from anatomically and electrophysiologically-identified PG cells (see above). As expected, one-second light pulses (590 nm LED) applied to the glomerulus elicited an 8 ± 2 mV hyperpolarization in PG cells (*p* = 0.02 in paired t-test, *n* = 4; **Supplementary Fig. 4B**) and nearly eliminated spontaneous or evoked spikes in these cells (*n* = 3; **Supplementary Fig. 4C**). In contrast, direct NpHR3-mediated currents were at most minimal in 6 eTC recordings. In addition, in GAD65-Cre-tdTomato mice, compressed z-stack confocal images displayed moderate fluorescence signals in small cell bodies that surrounded glomeruli, likely PG cells (**Supplementary Fig. 4D**). In some eTC recordings in GAD65-Cre-NpHR3 mice, including the example in **Supplementary Fig. 4B**, light appeared to evoke a very small inward current, but this likely reflected electrical coupling of signals generated in PG cells. In one eTC-PG cell pair recording, clear evidence for electrical coupling was observed when the test eTC was hyperpolarized with direct current injection.

A second issue in these experiments was the possibility that the eTC currents at the +28 mV holding potential used to isolate inhibition at least partially reflected glutamatergic currents with reversed polarity. The eTC recordings were conducted with cesium-containing electrodes (see above), heightening this possibility. That the currents were glutamatergic however was excluded by comparing the time courses of the eTC currents at +28 mV and –77 mV just after OSN stimulation (**Supplementary Fig. 4E**). During the time-window at which the OSN-EPSC peaked at –77 mV (∼3 ms after OSN stimulation), the currents at +28 mV were either near zero or small inward deflections. If the more delayed outward currents in the eTC at +28 mV were glutamatergic, the OSN-EPSC should have appeared as an outward current at +28 mV. The GABAergic nature of the outward currents near +28 mV was further confirmed by the fact that they were blocked by the GABA_A_ receptor blocker gabazine (Zak et al., 2015).

Finally, to further assess whether the OSN stimuli used in the eTC recordings mainly excited neurons at one glomerulus (McGann et al., 2005; Gire and Schoppa, 2009), we used a Förster energy resonance transfer (FRET)-based protein sensor of glutamate, FLII^81^E-1μ (Dulla et al., 2008; **Supplementary Fig. 5**), which reduces fluorescence upon binding of glutamate. After dye-loading (concentration = 50 ng/ml) in rat bulb slices, FRET signals (CFP excitation at ∼440 nm; Venus (Y) emission at ∼535 nm) were visualized by whole field epi-illumination using a Sutter Lambda DG-4 light source and the Zeiss Axioskop2 FS Plus slice microscope. FRET signals were recorded in the superficial 50 microns of the slice (100-200 images collected at 7-8 Hz, 50 msec exposures using SlideBook; Intelligent Imaging Innovations, Inc). Electrical stimulation of OSNs at low intensities (15-50 μA, 100 μsec) resulted in a decrease in the FRET signal within a target glomerulus (mean ΔF/F = –1.8 ± 0.4%; *n* = 4) that persisted for ∼0.5 seconds (**Supplementary Fig. 5A**). Neighboring glomeruli displayed much smaller signals (mean size relative to target glomerulus = 0.14 ± 0.03, 8 neighboring glomeruli across 4 experiments; **Supplementary Fig. 5A,C**), consistent with excitation being mainly limited to the target glomerulus. It is possible that the small fluorescence decreases observed in neighboring glomeruli were due to inadvertent capturing of fluorescence changes that originated in the target glomerulus. Indeed, in 3 experiments in which FLII^81^E-1μ–mediated signals were measured in glomeruli following direct depolarization of an eTC with a patch pipette (50 ms pulses to 0 mV), when evoked glutamate transients should have been restricted to target glomeruli, small decreases in fluorescence in neighboring glomeruli were also observed (mean size relative to target glomerulus = 0.12 ± 0.04, 5 neighboring glomeruli; **Supplementary Fig. 5B,C**). Validating that the FLII^81^E-1μ–mediated signals reflected glutamate transients, we found that application of the glutamate uptake blocker DL-TBOA (25 μM) prolonged the fluorescence transients (increase in half-width from 605 ± 25 ms under control conditions, *n* = 47 glomeruli, to 1080 ± 120 ms, *n* = 6 glomeruli; *p* < 0.01 in unpaired t-test).

### Statistics and analysis of electrophysiological data

Data were analyzed using Axograph or Microsoft Excel (Microsoft) and are generally expressed as mean ± SEM (except where noted). Significance was most commonly determined using the two-tailed Wilcoxon matched-pairs signed-rank test or the two-tailed Mann-Whitney U test. In a few matched pair comparisons with a smaller sample size (*n* = 4 or 5), a paired t-test was used. A value of *p* < 0.05 was considered significant (asterisks in the figures), except if multiple comparisons were being made, in which case the Bonferroni correction was applied. For assessing the effect of DL-TBOA on *EC_extra_* and *IC* (**Fig. 9**), Analysis of Covariance (ANCOVA) was used, wherein the associated conductances under control and DL-TBOA conditions were related to the amplitude of the OSN-EPSC.

Stimulus artifacts in many of the illustrated traces have been blanked or truncated. In the figure summary plots that reflect measurements of inward glutamatergic currents, values for current amplitude and integrated charge were multiplied by –1 to facilitate interpretation.

In the MC-PG cell pair recordings used for determining the mechanisms of eTC-to-MC extrasynaptic transmission (**Fig. 1**), an event detection routine was used to detect rapid EPSCs (or mEPSCs in the presence of TTx) in the PG cell and the co-occurring current in the MC was also captured. Analysis of PG cell EPSCs was conducted both on spontaneous activity (*n* = 3) or, more commonly, on current events evoked by OSN stimulation (*n* = 5). OSN stimulation elicited prolonged bouts of synaptic activity in PG cells that lasted seconds, reflecting sustained activation of the glomerular circuitry (Schoppa and Westbrook, 2001; Yuan and Knopfel, 2006; De Saint Jan and Westbrook, 2007). Because the analysis required precise temporal information about the timing of the currents in the PG cells versus MCs, EPSCs in the PG cells were selected for analysis that were separated from other EPSCs by at least 50 ms; the EPSC bursts that identified the PG cell as being the subtype that receives input from eTCs (see above) were avoided. In the triple-cell recordings, isolated action potential currents (> 50 ms-separation from other spikes) were detected in the eTC using a thresholding method, and currents locked to these spikes were averaged in the PG cell and MC. In the eTC-MC pair recordings used to test for spillover at OSN-to-eTC synapses (**Fig. 3**), a similar analysis to that used with the PG-MC pairs was performed, except rapid, spontaneously-occurring EPSCs in the eTC were first detected.

In the PG cell-MC pair recordings and eTC-PG cell-MC triple-cell recordings, some care was taken to ensure that the rapid EPSCs in the PG cell that were analyzed reflected eTC inputs and not alternate sources. For example, in experiments employing OSN stimulation, the first 50 ms after the stimulus was excluded in the analysis in order to avoid EPSCs that could have reflected monosynaptic OSN-to-PG cell inputs. Potential EPSCs in the PG cell that reflected MC-to-PG cell inputs were avoided using two criteria. First, by excluding the first 50 ms after the OSN stimulation in the analysis (in the OSN stimulation experiments), we limited the contribution of MC-to-PG cell inputs induced by spikes directly evoked by monosynaptic OSN-to-MC inputs (Najac et al., 2011). Also, more delayed MC-to-PG cell events were excluded by avoiding LLD events (see above) in the analysis. Prior slice studies have established that delayed spiking in MCs (and thus glutamate release) requires that a glomerulus engage in an LLD (Gire and Schoppa, 2009).

In the pair-cell recordings used to analyze the relationship between eTC spike number and *EC_extra_* in MCs/eTCs (**Fig. 4**), current integrals were measured over varying time-windows (50-700 ms) reflecting differences in the apparent duration of *EC_extra_* for different numbers of eTC spikes (see, for example, **Fig. 4A2**). From the measurements of charge associated with *EC_extra_*, we computed a parameter *S* that reflected the deviation from linearity in the rise of the current for increasing spike number (*N_spike_*) from:
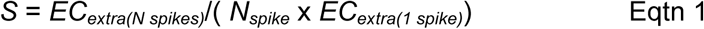

In this expression, *EC_extra(N spikes)_* was the magnitude of the extrasynaptic current in the MC (or eTC) evoked when the spiking eTC engaged in *N* spikes, while *EC_extra_*_(*1 spike*)_ was the extrasynaptic current evoked when the same eTC spiked once. Linearity, corresponding to when *S* = 1, occurred when the observed extrasynaptic current was the linear summation of the current evoked by one spike. *EC_extra_* in the pair-cell recordings was evoked by either direct depolarizing current injections in the eTC (while in whole-cell patch mode) or spontaneously-occurring spikes in the eTC (while in LCA patch mode). The results from the two types of experiments were similar (*p* = 0.28 in extracted values for *S*) and pooled. For the recordings from eTC-PG cell pairs (**Fig. 5**), a similar analysis was performed based on integrating the bursts of rapid EPSCs in the PG cell for eTCs engaged in *N* spikes. In the assessment of whether PG cells displayed slow, extrasynaptic currents (**Fig. 5E**), spontaneous EPSC bursts in PG cells that displayed ≥4 rapid EPSCs were selected for each recording, aligned to the start of the first EPSC in each burst, and then averaged. Currents were then integrated in a 100-ms window, 100-200 ms after the start of the first EPSC. In the initial selection of the rapid EPSC bursts prior to averaging, those that persisted for longer than 100 ms were excluded.

In the analysis of the eTC-PG cell pair recordings used to test whether recurrent excitation occurred during eTC spike bursts (**Fig. 7**), action potential currents in the eTC were first detected using a thresholding method and 16-ms intervals of PG cell current centered on the time of each eTC spike were selected. Rapid EPSCs were then detected from the 16-ms current intervals using an amplitude threshold on filtered (600-800 Hz) derivatives of the current. The threshold was selected based on visual inspection of whether the procedure effectively detected EPSCs in the PG cell with no false positives. Detected EPSCs that occurred in a window 0.5-3.5 ms after the eTC spike were considered to reflect direct transmission from the test eTC to the test PG cell; all other detected EPSCs were considered to reflect transmission from eTCs other than the test eTC (i.e., recurrent excitation). Because in our primary analysis we wished to exclude eTC spike bursts that were associated with the network-wide LLD, eTC spike bursts were initially sorted by whether there were co-occurring LLDs. In 3 of the 5 eTC-PG cell pairs, direct information was available about the LLD from either a simultaneous MC recording (in two triple-cell recordings) or from the eTC recording itself (in one case, when the eTC recording was in current-clamp mode). In the two other eTC-PG cell recordings, whether an eTC spike burst was associated with an LLD was inferred based on the analysis of the three experiments in which definitive information about the LLD was available. These results (summarized in **Fig. 7D**) indicated an unambiguous separation in the number of EPSCs in the PG cell that were not time-locked to spikes in the test eTC depending on whether the eTC spike burst was associated with an LLD.

In the analysis of excitatory currents evoked by OSN stimulation in eTCs (**Figs. 8,9**), we extracted measures of the isolated *EC_extra_* from the composite current by subtracting an estimate of the OSN-EPSC, based on a double-exponential fit of the rise and decay of the fast current component (**Fig. 8B2**). This method relied on the OSN-EPSC being discernible from *EC_extra_*, which was generally the case (see example traces in **Figs. 8** and **9**). Further supporting the validity of the method, we found that, for most stimuli, the estimated decay time-constant of the OSN-EPSC did not vary significantly with current amplitude (9 ± 5% increase in decay time constant for 4.9 ± 0.5-fold increase in OSN-EPSC amplitude, *n* = 4 eTC cells). Because *EC_extra_* became faster and was more likely to blend in with the OSN-EPSC with stronger stimuli (see example traces in **Fig. 8B1**), the decay constant should have increased with current amplitude if the fitted currents had included a significant *EC_extra_* component (since *EC_extra_* was still slower than the OSN-EPSC). When the OSN-EPSC was > 1 nA, *EC_extra_* and the OSN-EPSC were sometimes not discernible by eye; also, the currents were fitted with decay constants that were as much as 70% slower than that used to fit discernible OSN-EPSCs at lower stimulation intensities in the same cell. In these cases, the subtraction procedure for isolating *EC_extra_* likely led to a measurement that was smaller than its actual value. Thus, our analysis likely understated the extent to which the E/I balance favors excitation at high OSN input levels.

### Ultrastructural studies

The tissue sections from the olfactory bulb used for electron microscopy analysis were the same as those from Bourne and Schoppa (2017). The methods of preparing olfactory bulb slices used for filling eTCs with biocytin, tissue fixation, DAB-labeling, and the final slicing of thin (50 nm) sections prior to EM analysis, are described in that study. Sections were imaged either on a FEI Tecnai G2 transmission electron microscope at 80 kV with a Gatan UltraScan 1000 digital camera at a magnification of 4,800x or a Zeiss SUPRA 40 field-emission scanning electron microscope (FE-SEM) equipped with an integrated module called ATLAS™ (Automated Large Area Scanning; software version 3.5.2.385; Kuwajima et al., 2013).

The serial section images were aligned and dendrites, axons, and glia were traced using the RECONSTRUCT^TM^ software [available for free download at http://synapses.clm.utexas.edu] (Fiala and Harris, 2001; Fiala, 2005). For this study, we analyzed the neuropil surrounding dendrodendritic synapses between eTC dendrites and putative inhibitory dendrites as well as OSN synapses onto eTC dendrites. Although the dark precipitate formed by the DAB reaction obscures the interior of the eTC dendrite, presynaptic vesicles could still be identified and all analyzed dendrodendritic synapses had a clear asymmetric postsynaptic density (see insets of **Fig. 2A1** and **B1** for examples). Similarly, OSN synapses onto DAB-labeled eTC dendrites were identified based on docked presynaptic vesicles, a clear synaptic cleft, and often a visible asymmetric postsynaptic density (see inset of **Fig. 2C1** for example). Identification of the surrounding elements of the neuropil was based on their ultrastructural features. Glial processes were identified based on dark cytoplasm (likely due to endogenous peroxidase activity that reacted with the DAB – see Schipper et al., 1990), absence of vesicles, and long, thin morphology (Ventura and Harris, 1999). Excitatory dendrites had round, clear vesicles and formed asymmetric synapses. Inhibitory dendrites had flattened, pleiomorphic vesicles and formed symmetric synapses. Dendrites were distinguished from axons based on their lack of boutons, fewer vesicles, and the presence of efferent and afferent synapses.

Each of the eTC-to-PG cell synapses (13) and OSN-to-eTC synapses (39) was examined through serial sections to determine if glial processes or putative MC dendrites were nearby, within 0.5 μm. This number, 0.5 μm, was chosen to be in the range of extrasynaptic transmission in part based on the modeling studies of Pendyam and co-workers (2009), which showed that large supralinear increases in glutamate concentration could occur at distances from glutamate release sites near this value. Dendritic processes that formed at least one gap junction with another dendrite were assigned to be putative MC dendrites. This was based on evidence that the MC-to-MC gap junctional conductance is much greater than the eTC-to-MC or eTC-to-eTC gap junction conductance (Hayar et al., 2005; Gire et al., 2012). Any uncertainty that the putative MC dendrites actually reflected MCs (versus eTCs or other tufted cell sub-types) did not detract significantly from our conclusions about the mechanisms of extrasynaptic transmission. Our electrophysiological results indicated that eTCs send extrasynaptic signals not just to MCs but also other eTCs (**Fig. 4F**).

A subset (5) of the eTC-to-PG cell synapses were selected for a brick analysis to further characterize the surrounding neuropil (two examples in **Fig 2A,B**). A 2×2 micron grid was positioned with the synapse in the middle. Depending on the size of the synapse, the grid was copied to the same location on serial sections above and below the synapse to sample the surrounding neuropil, for a total of 14-20 sections (50 nm/section). Each structure (astroglia, putative excitatory dendrite, putative inhibitory dendrite, axon) contained within the brick was identified by following it through serial sections.

## Acknowledgements

The authors thank members of the Schoppa lab for helpful discussions.

---

**Supplementary Figure 1.**
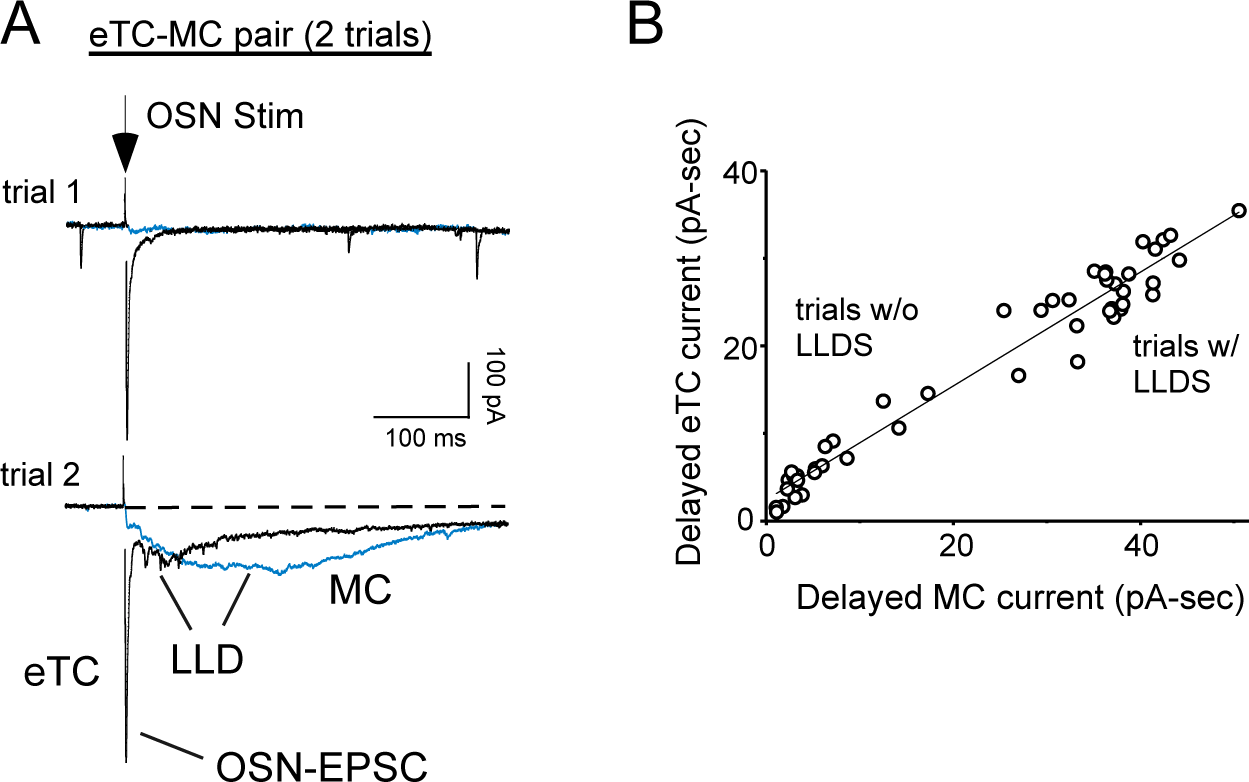
Physiological evidence that eTC-MC pairs are affiliated with the same glomerulus. **(A)** Example responses to OSN stimulation (38 μA) in one of the eTC-MC pair-cell recording (same as in **Fig. 2B**) used to test for spillover at OSN-to-eTC synapses. Two response trials overlaid for both cells are shown. Note that in trial 2, both cells displayed a delayed LLD current (Carlson et al., 2000) that was not observed in trial 1. **(B)** Summary plot generated from 49 response trials from the experiment in **A.** Note the high degree of correlation in the delayed current magnitude in the two cells (linear regression fit: *r* = 0.98, *p* < 1,0e-06) and co-occurrence of the LLDs. Current magnitude measurements were obtained by integrating the current starting 50 ms after OSN stimulation in order to avoid the OSN-EPSC.

**Supplementary Figure 2.**
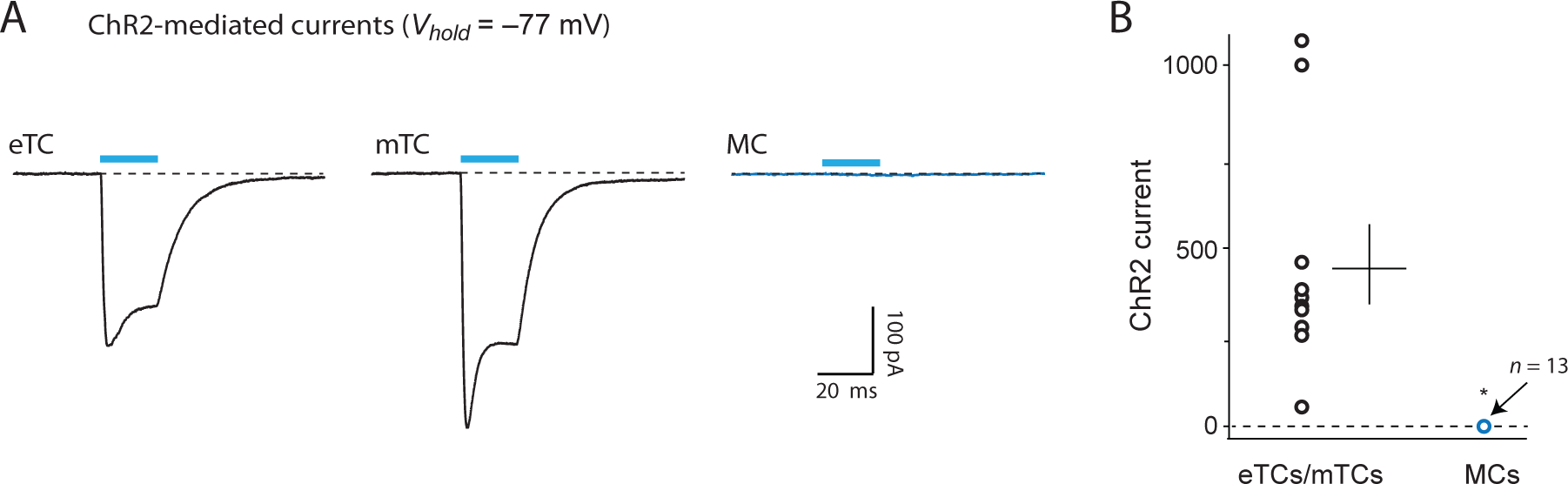
Tests for direct light-evoked currents in CCK-Cre-ChR2 mice. (**A**) Example recordings of light-evoked (20 ms pulse; 473 nm LED) ChR2-mediated currents in an eTC (left) and a middle TC (mTC; middle), and the absence of such a current in a MC (right). *EC_extra_* was not present in the MC response because recordings were condicted in the presence of NBQX (10 μM) and DL-AP5 (50 μM). Traces reflect averages of at least 8 trials. (**B**) Summary plot illustrating ChR-mediated current amplitude measurements from tufted cells (*n* = 10; 2 eTCs and 8 mTCs) and MCs (*n* = 13). For the tufted cells, lines reflect mean ± SE of the peak inward current. Values were multipled by –1. For MCs, values ranged from –3 pA to +3 pA. \**p* < 0.01 in Mann-Whitney U test.

**Supplementary Figure 3.**
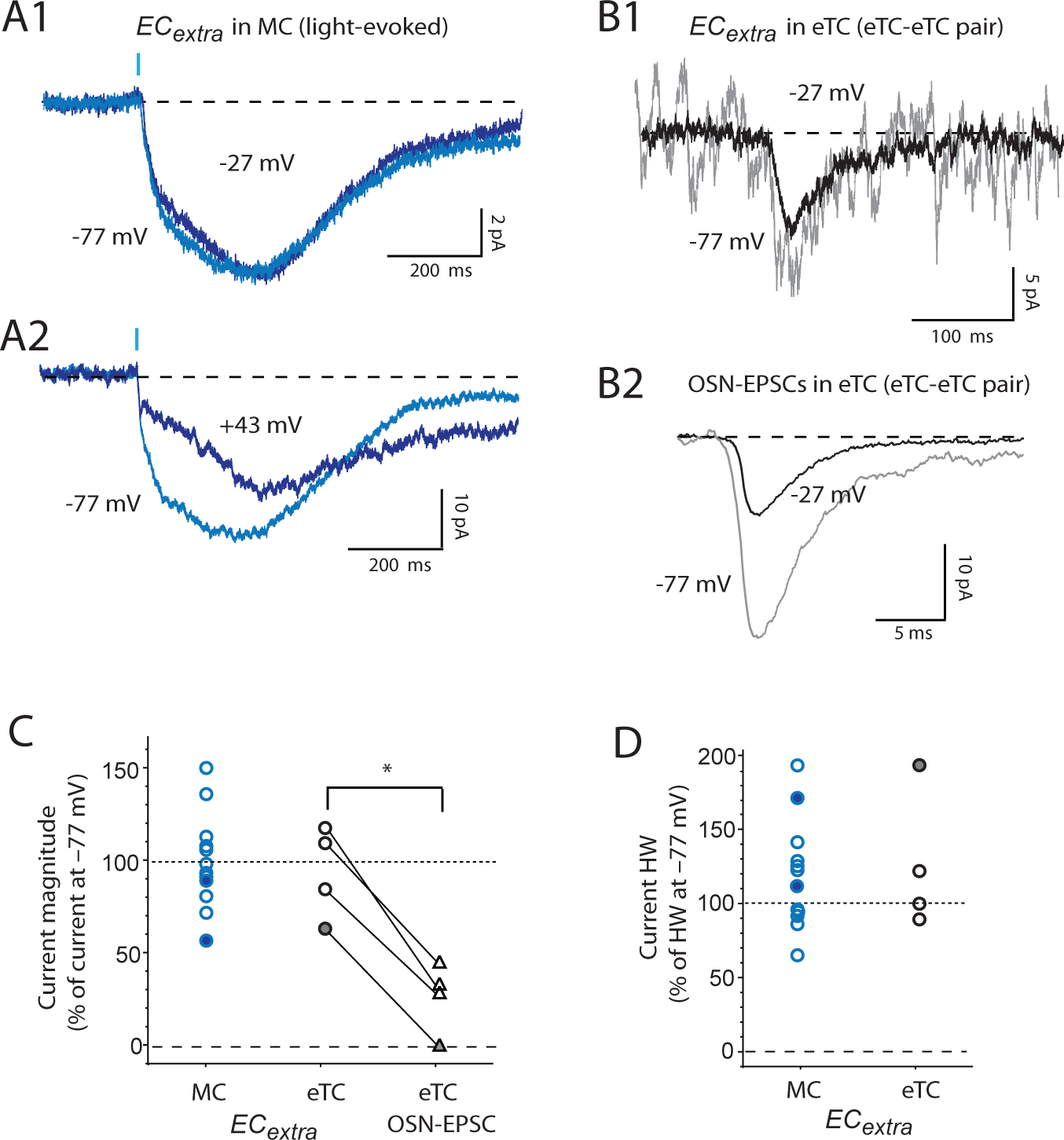
Voltage-dependence of *EC_extra_*. (**A**) Example recordings of light-evoked (0.1 ms pulse; 473 nm LED) *EC_extra_* in MCs in CCK-Cre-ChR mice, comparing responses in the same cell at depolarized voltages (–27 mV in **A1**; +43 mV in **A2**) versus the standard hyperpolarized voltage (–77 mV). Note that depolarization caused at most only modest changes in the magnitude and kinetics of *EC_extra_*. Traces reflect averages of 11-33 responses. (**B**) Recordings of *EC_extra_* (**B1**; averages of 10-33 trials) and spontaneous OSN-EPSCs (**B2**; averages of 68-82 detected events) at different voltages in the same eTC during an eTC-eTC pair recording. Depolarizing the holding potential to –27 mV did not significantly impact *EC_extra_* (induced by 50 ms depolarization to other eTC, not shown) versus that at –77 mV, but did greatly reduce the OSN-EPSC. (**C**) Summary of the effect of depolarizing the membrane potential on the magnitude of *EC_extra_* in MCs (blue symbols; light evoked) and eTCs (black symbols; depolarization-evoked in eTC-eTC pairs). *EC_extra_* magnitudes at depolarized holding potentials (open symbols = –27 mV; filled symbols = +28 or +43 mV) are plotted relative to magnitude at –77 mV in the same cell. Recordings of OSN-EPSCs in eTC-eTC pairs are included for comparison. \**p* = 0.0025 in paired t-test comparing effect of depolarization on *EC_extra_* versus OSN-EPSCs in eTC-eTC pairs. (**D**) Summary of the effect of depolarization on *EC_extra_* half-width (HW). Same *EC_extra_* recordings as in **C**.

**Supplementary Figure 4.**
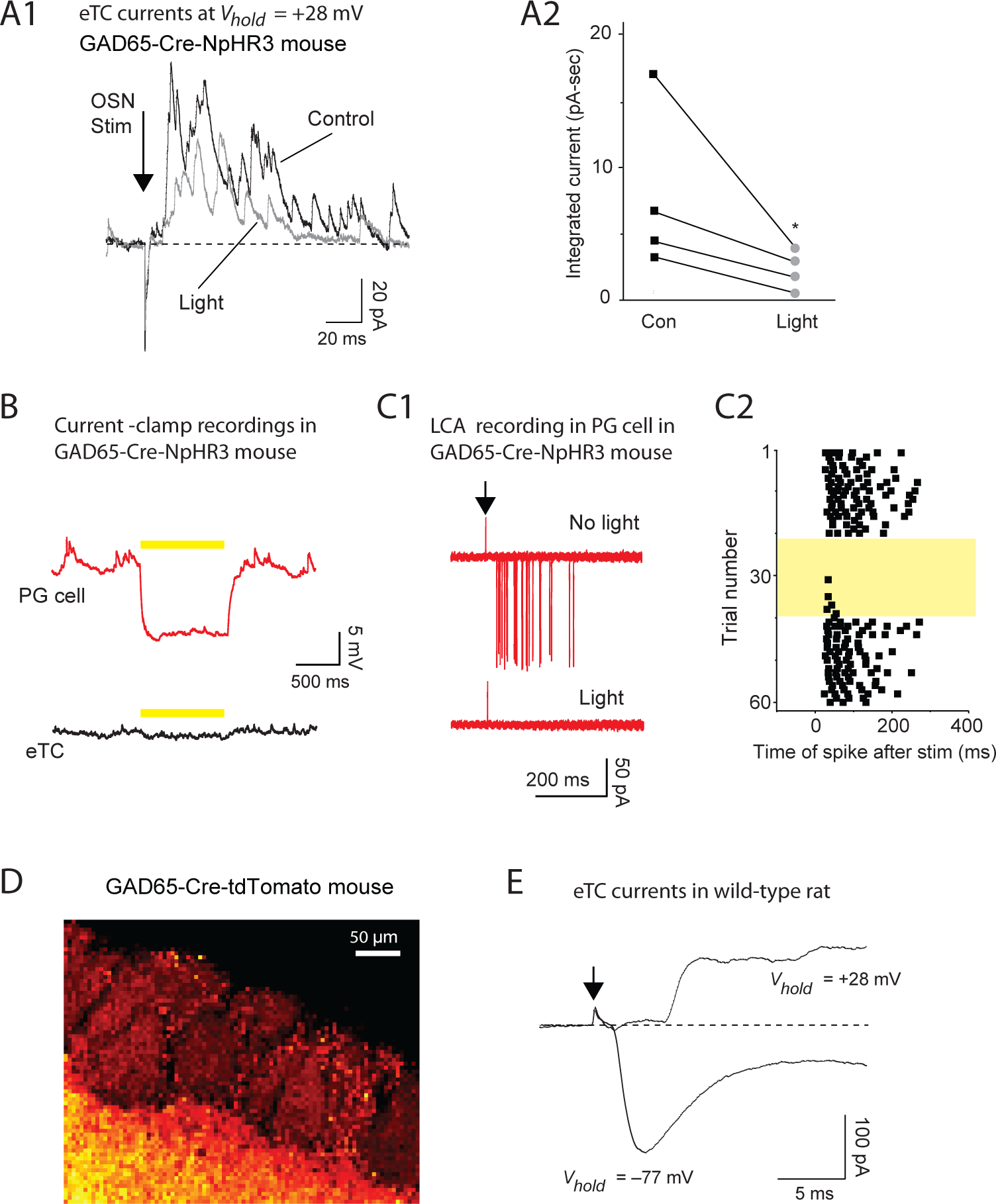
Analysis of outward currents in eTCs in response to OSN stimulation. (**A1**) Inhibitory currents in an eTC evoked by OSN stimulation (18 μA) in a GAD65-Cre-NpHR3 mouse, showing reduction of the current by 590 nm LED light (320 ms pulse, starting 20 ms prior to OSN stimulation). Data traces reflect averages of 5 trials. (**A2**) Summary of 4 experiments similar to **A1**. \**p* = 0.004 in paired t-test conducted on normalized data. (**B**) A 1-second light pulse (590 nm LED) in a GAD65-Cre-NpHR3 mouse elicited a hyperpolarization in a PG cell (top), as expected (Kiyokage et al., 2010), but a negligible signal in an eTC. The apparent very small hyperpolarization in the eTC likely reflected electrical coupling of signals originating in the PG cell (see *Methods*). Traces reflect averages of 5-7 trials. (**C**) Light pulses in GAD65-Cre-NpHR3 mice also reduced spiking in a PG cell evoked by electrical stimulation of OSNs (8 μA). Raw traces (**C1**; 5 superimposed trials for each condition) and the time-course of spiking across trials (**C2**; each square = 1 spike) are illustrated. Yellow denotes period in which LED pulses were applied. **(D)** Fluorescence measurement (compressed z-stack confocal image) from an olfactory bulb slice taken from a GAD65-Cre-tdTomato mouse. Note the moderate signal in the small cells surrounding glomeruli, likely PG cells. The bright diffuse signal in the EPL likely reflected the dendrites of granule cells. (**E**) Comparison of the early component of the eTC current response to OSN stimulation recorded at +28 mV and –77 mV (same 3-μA current trace as in **Fig. 8B1** but expanded; averages of 5 trials). Note that, at +28 mV, there is near-zero current just after OSN stimulation, corresponding to when the OSN-EPSC at –77 mV peaked. This indicated that +28 mV was near the reversal potential for excitatory synaptic currents and that the more delayed outward current did not reflect glutamatergic currents of reversed polarity.

**Supplementary Figure 5.**
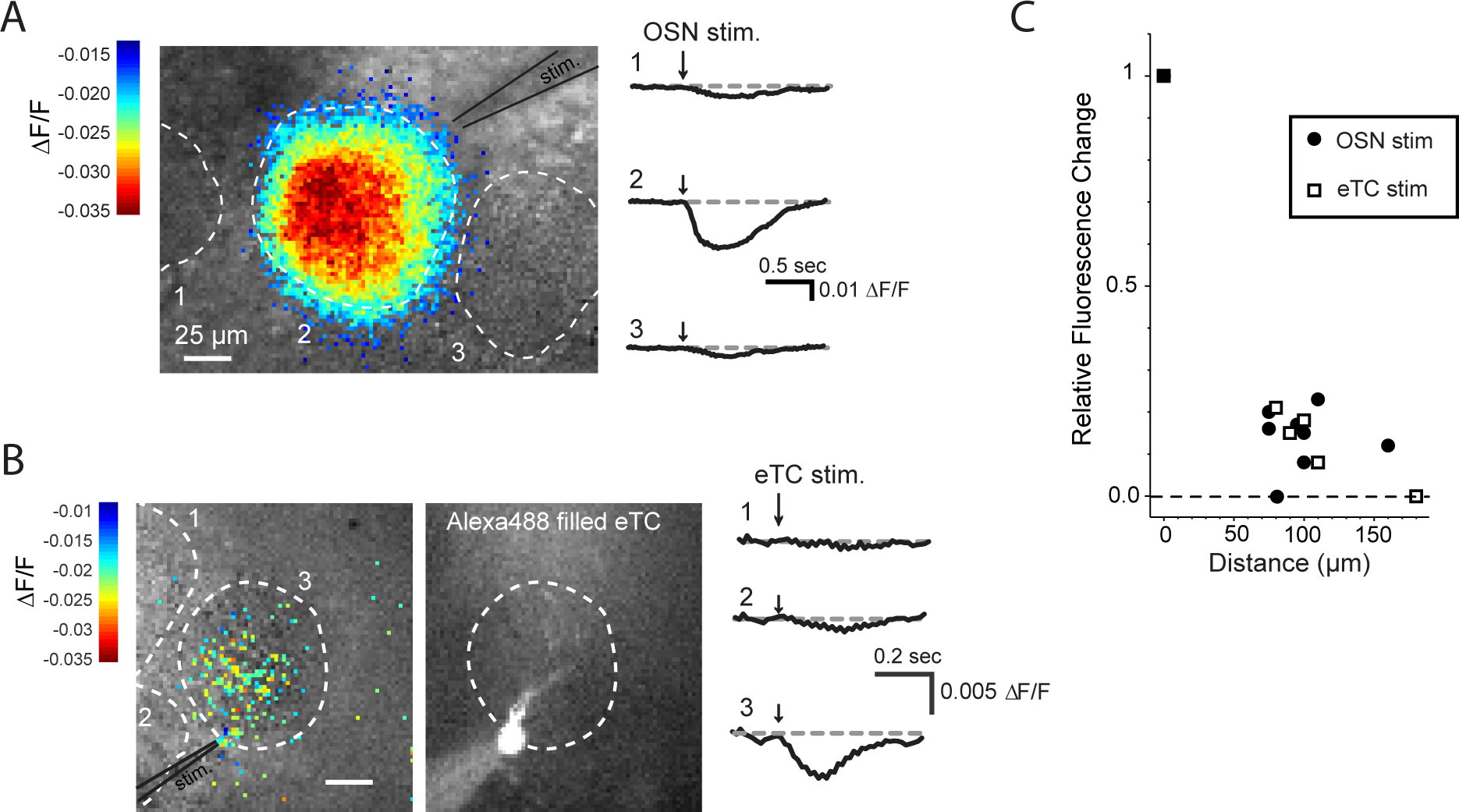
Recordings of glutamate transients in glomeruli with a FRET-based biosensor. (**A**) Example FLII^81^E-1μ-mediated fluorescence responses (YFP signals at ∼535 nm) following electrical stimulation of OSN axons (15 μA; stimulating electrode indicated). Note the large decrease in fluorescence at the glomerulus nearest the stimulating electrode, observed both in the image (*left*) and time-courses (*right*). Neighboring glomeruli displayed much smaller responses, consistent with stimulation mainly exciting neurons at one glomerulus. (**B**) A different example of fluorescence responses evoked by direct depolarization of an eTC with a whole-cell patch electrode (50 ms pulse from –70 mV to 0 mV). The image at left and time courses at right show that the signals were largely restricted to the target glomerulus of the eTC, as expected for this stimulation method. The image in middle illustrates the test eTC filled with Alexa 488. (**C**) Summary of fluorescence measurements across 4 OSN stimulation experiments (filled circles) and 3 eTC stimulation experiments (open squares). Each data point reflects the fluorescence change at one glomerulus that was near the target glomeruli. Results are plotted as the change in fluorescence relative to that observed in the target glomerulus versus the distance between the test and target glomeruli (measured from the glomerular centers).

**Supplementary Figure 6.**
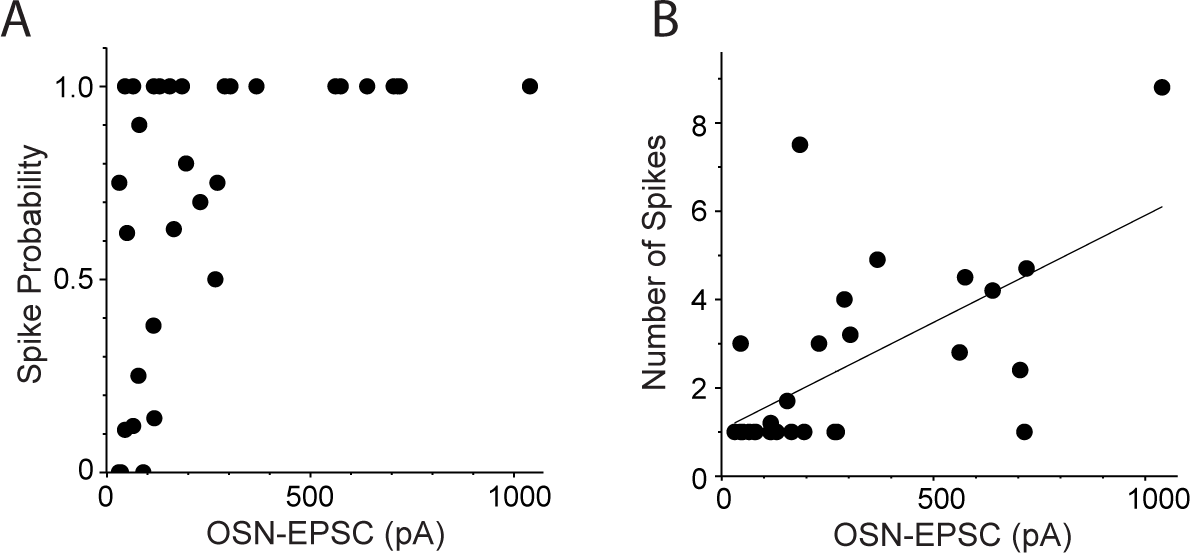
The number of spikes evoked in eTCs by OSN stimulation is correlated with the OSN-EPSC amplitude. (**A**) Relationship between the peak amplitude of the OSN-EPSC and the probability of spiking in eTCs. Data points in each plot reflect 16 eTC whole-cell recordings that transitioned between voltage-clamp and current-clamp to measure both OSN-EPSCs and spiking. Some experiments produced multiple data points reflecting the fact that multiple OSN stimulus intensities were sampled. (**B**) Relationship between the peak amplitude of the OSN-EPSC and the number of evoked spikes in eTCs, excluding failures (trials with no spikes). Data were fitted to a line to estimate a correlation coefficient (*r* = 0.62, *p* = 0.0032, *n* = 29). Same experiments as in **A**.

